# Improved dual-color GRAB sensors for monitoring dopaminergic activity *in vivo*

**DOI:** 10.1101/2023.08.24.554559

**Authors:** Yizhou Zhuo, Bin Luo, Xinyang Yi, Hui Dong, Jinxia Wan, Ruyi Cai, John T. Williams, Tongrui Qian, Malcolm G. Campbell, Xiaolei Miao, Bozhi Li, Yu Wei, Guochuan Li, Huan Wang, Yu Zheng, Mitsuko Watabe-Uchida, Yulong Li

## Abstract

Dopamine (DA) plays multiple roles in a wide range of physiological and pathological processes via a vast network of dopaminergic projections. To fully dissect the spatiotemporal dynamics of DA release in both dense and sparsely innervated brain regions, we developed a series of green and red fluorescent GPCR activation‒based DA (GRAB_DA_) sensors using a variety of DA receptor subtypes. These sensors have high sensitivity, selectivity, and signal-to-noise properties with subsecond response kinetics and the ability to detect a wide range of DA concentrations. We then used these sensors in freely moving mice to measure both optogenetically evoked and behaviorally relevant DA release while measuring neurochemical signaling in the nucleus accumbens, amygdala, and cortex. Using these sensors, we also detected spatially resolved heterogeneous cortical DA release in mice performing various behaviors. These next-generation GRAB_DA_ sensors provide a robust set of tools for imaging dopaminergic activity under a variety of physiological and pathological conditions.

## Main

Dopamine (DA) is a physiologically essential monoamine neuromodulator released by dopaminergic neurons that project throughout the central nervous system. Interestingly, high spatial heterogeneity in terms of dopaminergic innervation—and therefore DA release—has been reported in various brain regions^1–3^. The DA system is known for its roles in reward and reinforcement learning, motor function^3,4^, memory consolidation^5,6^, and emotional control^7^. These processes are mediated by dopaminergic circuits originating from the midbrain projecting to the striatum and nucleus accumbens (NAc), with the dorsal striatum and NAc receiving dense dopaminergic innervation. In contrast, the medial prefrontal cortex (mPFC) and amygdala receive relatively sparse dopaminergic innervation that regulates a wide range of brain functions important in mediating cognitive function^8,9^, social interactions^10^, and aversive sensing^11,12^. However, due to limitations in our ability to detect DA with high sensitivity and resolution, the spatiotemporal dynamics of dopaminergic transmission in these sparsely innervated brain regions remain largely unknown, particularly under various *in vivo* conditions. The ability to directly visualize and compare the dynamics of DA release in both densely and sparsely innervated regions under behaviorally relevant conditions will therefore provide valuable information regarding the spatiotemporal regulation of dopaminergic activity in the brain. However, measuring DA with high sensitivity in order to understand how dopaminergic signaling is affected by neuronal activity and/or other neuromodulators requires multiplexed DA imaging combined with optogenetics and the simultaneous imaging of other neurochemical processes.

Recent advances in the development of genetically encoded fluorescent sensors for detecting DA have led to robust tools that can measure dopaminergic signals at high spatial and temporal resolution. By combining a G protein-coupled receptor (GPCR) with circularly permutated fluorescent protein (cpFPs), our group and Tian’s group developed a series of genetically encoded green and red fluorescent DA sensors called GRAB_DA_ and dLight^13–16^, which can be used to measure DA release under physiological and pathological conditions with high spatiotemporal resolution. These early generation of sensors enabled us to expand the knowledge of spatiotemporal dynamics of DA transmission in reward, learning and movement^17–20^, in strongly innervated regions such as striatum where dopamine levels are high. However, tracking slight changes in DA levels *in vivo*, including tonic DA release and fluctuations in sparsely innervated regions, was less possible because of limitations in sensitivity, especially with bulk measurements like photometry. In addition, even in dense areas, better sensors with improved sensitivity and signal-to-noise ratio are required for DA imaging with higher spatiotemporal precision. Moreover, the performance of previously developed red-shifted DA sensors is relatively poor compared to green fluorescent sensors, greatly limiting their use. To overcome these issues, we performed large-scale rational mutagenesis and cell-based screening in order to develop a next-generation series of red and green fluorescent GRAB_DA_ sensors with extremely high sensitivity, a high signal-to-noise ratio, and a wider concentration detection range for measuring DA release in a wide range of brain regions.

## RESULTS

### Engineering and characterization of DA sensors in cultured cells

To obtain next-generation DA sensors with improved sensitivity, selectivity, and distinct pharmacology profiles, we used various DA receptor subtypes cloned from several species as the sensor scaffold and replaced each receptor’s third intracellular loop (ICL3) with the ICL3 previously used in existing GRAB sensors^13,15,16,21^. We identified several chimera prototypes with promising performances in the initial screening by transplanting the ICL3 of existing sensors to different sites dopamine receptors, for example, transplanting the ICL3 of rGRAB_DA_ to Solenopsis invicta dopamine D2-like receptor (hereafter termed red fire ant D_2_R) and the ICL3 of GRAB_NE_ to bovine D_1_R. Interestingly, we also obtained good candidates when re-engineering the ICL3s of dLight1.3b^13^ and RdLight1^15^ to their original GPCR backbone, i.e., the human D_1_R (Fig. 1a; Extended Data Figs. 1 and 2). We then systematically optimized the length and amino acid composition of the linker sequences, key residues in the cpFP that affect protein folding and/or fluorescence intensity^22–27^, and sites in the GPCR that affect ligand binding and/or structural coupling^28–31^, and screened a total of 5000 variants (Extended Data Figs. 1 and 2). Using maximum brightness and the DA-induced change in fluorescence as our selection criteria, our screening yielded a series of top-performing DA sensors with various DA receptor backbones, including the green fluorescent gDA3m (based on human D_1_R) and gDA3h (based on bovine D_1_R) sensors, the red-shifted rDA2m and rDA2h (based on red fire ant D_2_R) sensors, and the red-shifted rDA3m and rDA3h (based on human D_1_R) sensors (Fig. 1b and 1c; Extended Data Fig. 4a), with “m” and “h” referring to medium and high DA affinity, respectively. We also generated DA-insensitive versions of these sensors by introducing mutations in the ligand-binding pocket of the corresponding GPCRs, yielding gDA3mut, rDA2mut, and rDA3mut for use as negative controls (Extended Data Figs. 1-3).

**Fig. 1.**
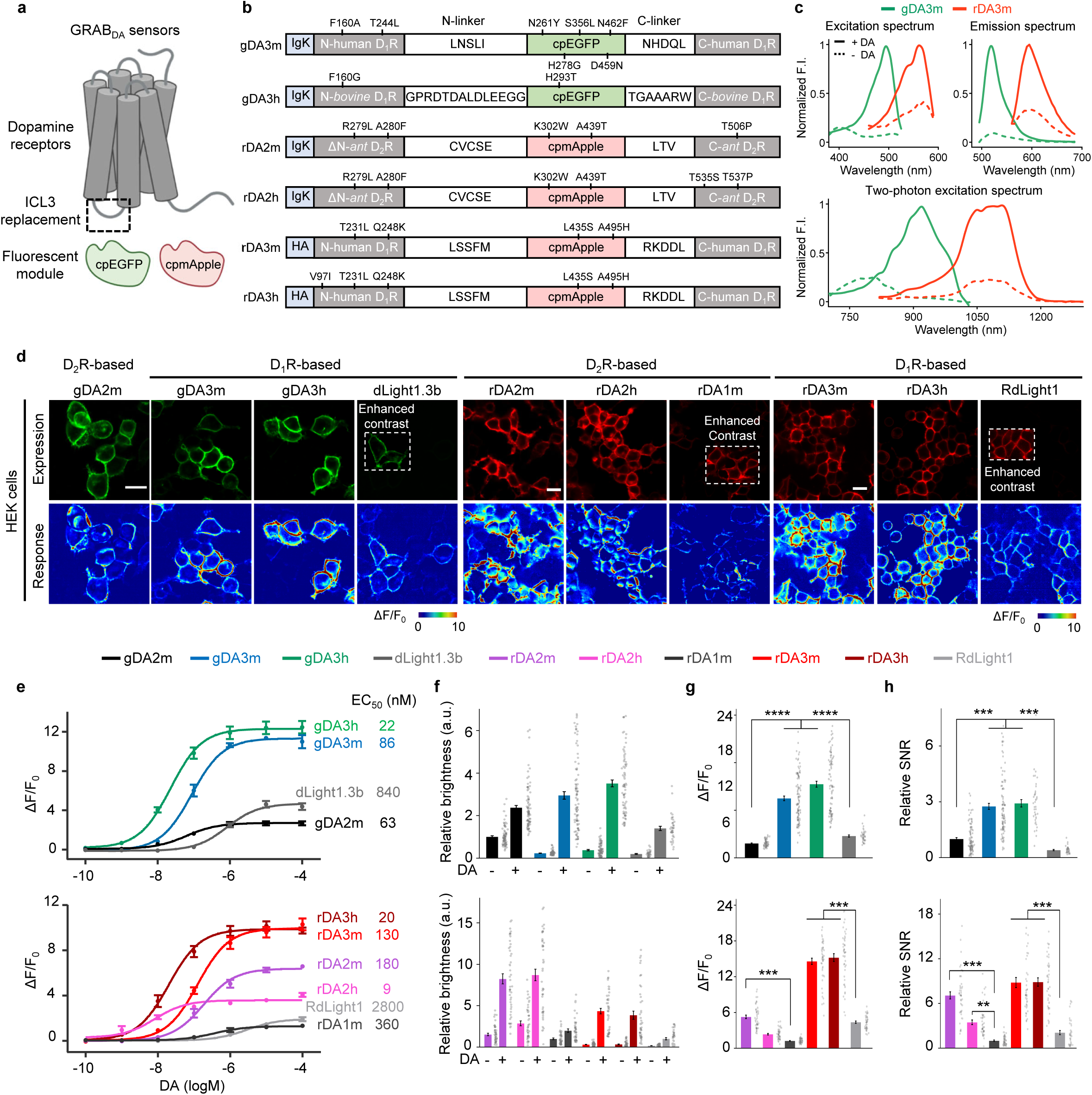
Development and performance of improved dual-color GRAB_DA_ sensors. **a**, Schematic illustration showing the principle of next-generation green and red fluorescent dopamine sensors. **b**, Schematics of improved dual-color GRAB_DA_ sensors. Mutations are indicated with respect to wild-type receptors and fluorescent proteins. Igk or HA, N terminus leader sequence. **c**, Spectral profiles of GRAB_DA_ sensors. One-photon excitation (top-left), emission (top-right) and two-photon excitation (bottom) spectra of indicated sensors in the absence (dashed lines) or presence (continuous lines) DA are shown. **d**, Representative images showing sensor expression (top) and fluorescence response to 100 μM DA (bottom) of indicated sensor variants. Scale bar, 20 μm. **e**, Titration DA curves of indicated sensors on HEK293T cells. Apparent affinity values (apparent EC_50_) are defined as the concentration of half-maximal fluorescence changes (max *ΔF/F_0_*). Data are shown as mean±SEM. *n*=4 wells with 400-500 cells per well for gDA3m, 7 for gDA3h, 10 for gDA2m, 8 for dLight1.3b, 7 for rDA2m, 7 for rDA2h, 6 for rDA1m, 12 for rDA3m, 12 for rDA3h, 12 for RdLight1. **f**, Group summary of relative brightness of indicated sensors before and after 100 μM DA addition. The brightness of green and red sensors was relative to DA2m and rDA1m, respectively. a.u., arbitrary unit. *n*=90 cells from 5 separate experiments (hereafter denoted as 90/5) for gDA3m, 82/5 for gDA3h, 88/5 for gDA2m, 37/3 for dLight1.3b, 45/3 for rDA2m, 45/3 for rDA2h, 45/3 for rDA1m, 45/3 for rDA3m, 45/3 for rDA3h, 38/3 for RdLight1. One-way ANOVA, post hoc Tukey’s test was performed. **g**, Group summary of maximal *ΔF/F_0_* of indicated sensors in response to 100μM DA. *n*= 130/7 for gDA3m, 173/8 for gDA3h, 200/10 for gDA2m, 80/4 for dLight1.3b, 45/3 for rDA2m, 45/3 for rDA2h, 45/3 for rDA1m, 45/3 for rDA3m, 45/3 for rDA3h, 38/3 for RdLight1. One-way ANOVA, post hoc Tukey’s test was performed. n.s. p=0.2820 between rDA2h and rDA1m. **h**, Group summary of signal-to-noise ratio (SNR) of indicated sensors. SNR was calculated based on the response to 100μM DA and was relative to DA2m (green) and rDA1m (red), respectively. *n*= 95/5 for gDA3m, 40/3 for gDA3h, 96/5 for DA2m, 31/3 for dLight1.3b, 45/3 for rDA2m, 45/3 for rDA2h, 45/3 for rDA1m, 45/3 for rDA3m, 45/3 for rDA3h, 38/3 for RdLight1. One-way ANOVA, post hoc Tukey’s test was performed. p=0.0016 between rDA2h and rDA1m.

All six of our newly generated GRAB_DA_ sensors localized well to the cell membrane when expressed in HEK293T cells and exhibited a large increase in fluorescence in response to bath application of 100 μM DA (Fig. 1d); moreover, the mutant versions were expressed at the cell surface but failed to respond to DA application (Extended Data Fig. 3). The sensors’ affinities were within physiological DA levels and ranged from nanomolar to submicromolar concentrations, with EC_50_ values of 22 nM and 86 nM for gDA3h and gDA3m, respectively, and 9 nM, 180 nM, 20 nM, and 130 nM for rDA2h, rDA2m, rDA3h, and rDA3m, respectively (Fig. 1e and Supplementary Table 1). We also compared these new sensors’ performance with previously reported GRAB_DA_ (gDA2m and rDA1m) and dLight (dLight1.3b and RdLight1) sensors^13–16^ in cultured cells. With respect to the green fluorescent sensors, both gDA3m and gDA3h had a >2-fold larger increase in fluorescence (with peak *ΔF/F_0_* values of ∼1000%) and a higher signal-to-noise ratio (SNR) compared to gDA2m and dLight1.3b (Fig. 1f-h and Supplementary Table 1). With respect to the red-shifted sensors, relative to rDA1m the basal fluorescence intensity values were 152%, 282%, 30%, 33%, and 16% for rDA2m, rDA2h, rDA3m, rDA3h, and RdLight1, respectively; moreover, rDA2m and rDA2h had the largest dynamic range (with peak *ΔF/F_0_* values of ∼560% and ∼240%, respectively) among the D_2_R-based red-shifted sensors (Fig. 1f-h and Supplementary Table 1). Finally, rDA3m and rDA3h had significantly higher brightness levels, fluorescence responses (with *ΔF/F_0_* values of ∼1000%), and SNR compared to RdLight1 (Fig. 1f-h). These results suggest that these next-generation GRAB_DA_ sensors might be useful for imaging DA release *in vivo* both in DA-abundant conditions and in brain regions with sparse dopaminergic innervation.

Next, we examined the properties of our new sensors when expressed in cultured neurons. Consistent with our results obtained using HEK293T cells, we found that the sensors localized well to the neuronal membrane (both at the cell body and in the surrounding neurites) and responded strongly to DA application (Fig. 2a-d), with DA affinity similar to what we measured in HEK293T cells. In addition, we obtained the same rank order in terms of the peak response to DA measured in cultured neurons and HEK293T cells. Moreover, the next-generation GRAB_DA_ sensors had higher SNR values compared to previous sensors when expressed in neurons (Fig. 2c and 2d and Supplementary Table 1).

**Fig. 2.**
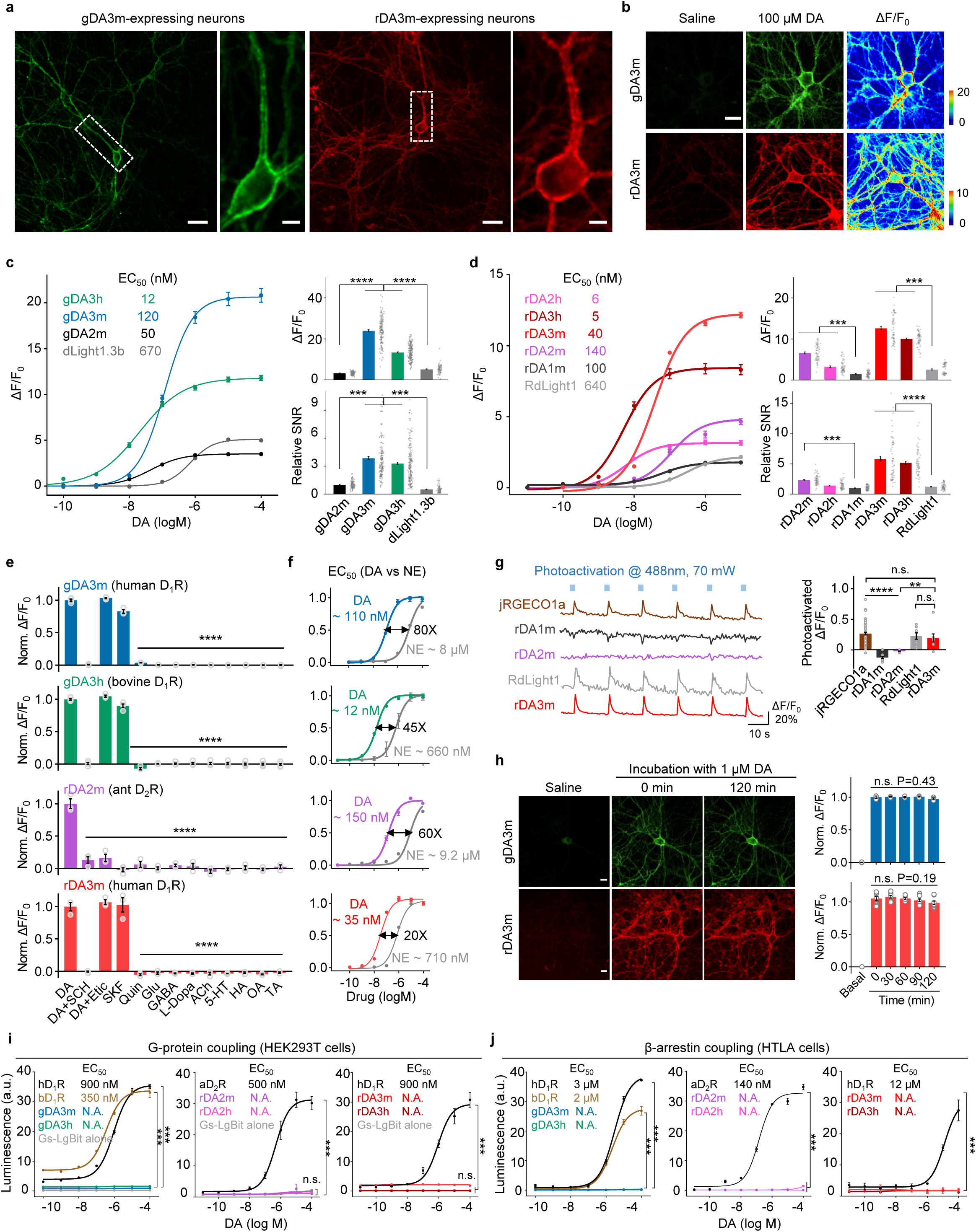
Characterization of new GRAB_DA_ sensors in cultured cells. **a**, Representative images of gDA3m or rDA3m expressing-primary cultured rat cortical neurons showing cell membrane localization. Scale bar, 100 μm (left) and 20 μm (right). **b**, Representative images showing expression and fluorescence response of gDA3m and of rDA3m in dissociated cortical neurons. Scale bar, 50 μm. Similar results were observed more than 30 cells. **c**, Titration DA curves (left) and group summary of peak response (top-right) and relative SNR (bottom-right) of indicated green dopamine sensors on cultured neurons. Left, *n*= 60, 60, 120, 60 neurons for gDA2m, gDA3m, gDA3h and dLight1.3b. Right, *n*=130, 175, 200, 80 neurons respectively. One-way ANOVA, post hoc Tukey’s test was performed. **d**, Titration DA curves (left) and group summary of peak response (top-right) and relative SNR (bottom-right) of indicated red dopamine sensors on cultured neurons. Left, *n*=30 neurons for all sensors. Right, *n*=60 neurons for all sensor variants. One-way ANOVA, post hoc Tukey’s test was performed. For SNR, n.s. p=0.7632 between rDA2h and rDA1m. ***p=0.005 between rDA2m and rDA1m. **e**, Pharmacological specificity of gDA3m, gDA3h, rDA2m and rDA3m in neurons. SCH-23390 (SCH), D_1_R antagonist; eticlopride (Etic), D_2_R antagonist; SKF-81297 (SKF), D_1_R agonist; quinpirole (Quin), D_2_R agonist; glutamate (Glu); gamma-aminobutyric acid (GABA); levodopa (L-Dopa); acetylcholine (ACh); serotonin (5-HT); histamine (HA); octopamine (OA); tyramine (TA). Antagonists were applied at 10μM, others at 1μM. *n*=4 wells for gDA3m, *n*=5 wells for gDA3h, *n*=3 wells for rDA2m and rDA3m. Each well contains 100-200 neurons. One-way Anova, post hoc Dunnett’s test was performed. gDA3m, n.s. p=0.0987 between DA and DA+Etic. gDA3h, n.s. p=0.2032 between DA and DA+Etic. rDA3m, n.s. p=0.8251, 0.9993 between DA and DA+Etic, or SKF. **f**, Titration curves of indicated sensors for the response to DA or NE in cultured neurons. Data are shown as mean±SEM. *n*=60 neurons from 3 experiments. **g**, Representative traces (left) and group summary of *ΔF/F_0_* (right) of indicated sensors upon blue-light illumination. *n*=70, 13, 17, 8, 8 cells for jRGECO1a, rDA1m, rDA2m, RdLight1 and rDA3m, respectively. One-way ANOVA, post hoc Tukey’s test was performed. n.s. p=0.7180 between jRGECO1a and rDA3m, p=0.9927 between rDA3m and RdLight1, **p=0.0083 between rDA2m and rDA3m. **h**, Representative images (left) and quantification (right) of the change in sensor fluorescence in response to 2-h application of 100μM DA. Scale bar, 20 μm. *n*=3, 9 cultures for gDA3m and rDA3m. One-way ANOVA test was performed for DA-containing groups. n.s. p=0.4375, 0.1895 for gDA3m, rDA3m. **i**, G-protein coupling was measured using luciferase complementation assay in cells expressing wild-type receptor, sensor or no receptor (G_s/i_-LgBit alone, ctrl). Data are shown as mean±SEM. *n*=3 cultures. **j**, β-arrestin coupling was measured with Tango assay in cells expressing receptor or sensor. Data are shown as mean±SEM. *n*=3 cultures. N.A., not applicable.

Importantly, our new sensors also retained the pharmacological specificity of their respective parent receptors. For example, application of the D_1_R-specific and D_2_R-specific antagonists SCH-23390 (SCH) and eticlopride (Etic), respectively, eliminated the corresponding sensors’ response to DA (Fig. 2e and Extended Data Fig. 5); interestingly, however, both the D_1_R-specific and D_2_R-specific antagonists inhibited the red fire ant D_2_R‒based rDA2m and rDA2h sensors (Fig. 2e and Extended Data Fig. 5b-d), possibly due to low sequence homology between red fire ant D_2_R and human D_2_R. Moreover, these new sensors had only a negligible response to a variety of other neurochemicals and transmitters, including glutamate, GABA, levodopa, acetylcholine, serotonin, histamine, octopamine, and tyramine. Importantly, despite the structural similarity between the transmitters DA and norepinephrine (NE) our optimized sensors were approximately 20-80-fold more sensitive to DA than NE (Fig. 2f and Extended Data Fig. 5), indicating their extremely high specificity for DA.

Next, we measured the kinetics of our DA sensors using rapid line-scanning confocal microscopy. We locally applied DA and then measured the time constant of the signal rise (τ_on_) and the time constant of the signal decay following application of the corresponding antagonist (τ_off_). Our analysis revealed τ_on_ values of approximately 80 ms for all DA sensors, and τ_off_ values ranging from 0.6-3 s based on differences in each sensor’s affinity (Extended Data Fig. 6 and Supplementary Table 1).

We then tested whether our red-shifted cpmApple-based DA sensors are photoactivated by blue light, as shown previously for the cpmApple-based red calcium indicator jRGECO1a^25,32^. Interestingly, unlike most mApple-based sensors, we found that bursts of 488-nm blue light had no significant effect on the fluorescence of the rDA1m and rDA2 sensors (Fig. 2g and Extended Data Fig. 4c and 4d and Supplementary Table 1), promising an optimal compatibility with blue-light-activated optogenetic actuators. However, though undesired, RdLight1 and rDA3 sensors were found to exhibit photoactivation when illuminated with blue light (similar as jRGECO1a), causing a transient increase in fluorescence (Fig. 2g and Extended Data Fig. 4c and 4d and Supplementary Table 1). In addition, the DA-induced increase in fluorescence was stable for up to 2 h in the continuous presence of 100 μM DA when expressed in cultured neurons, with minimal arrestin-mediated internalization or desensitization (Fig. 2h and Extended Data Fig. 4e-h). Thus, these sensors are suitable for long-term monitoring of dopaminergic activity.

To examine whether the DA sensors couple to intracellular signaling pathways, we used the luciferase complementation assay^33^ and the Tango assay^34^ to measure G protein‒mediated signaling and β-arrestin signaling, respectively. We found that wild-type receptors showed robust coupling in both assays, whereas all of the DA sensors tested failed to engage either of these GPCR-mediated downstream pathways (Fig. 2i and 2j). We therefore conclude that expressing these receptors likely does not affect cellular physiology.

### Imaging DA dynamics in acute brain slices

Next, we used two-photon imaging to measure the sensitivity of the gDA3m and rDA3m sensors for reporting the triggered release of endogenous DA in acute brain slices. We injected the nucleus accumbens (NAc)—which receives dense innervation from midbrain dopaminergic neurons (DANs)— with adeno-associated virus (AAV) expressing either gDA3m or rDA3m and then prepared acute brain slices two weeks after injection (Fig. 3a and 3b). Electrical stimuli applied at NAc at 20 Hz induced robust transient increases in fluorescence, with the magnitude of the peak response increasing with increasing numbers of stimuli. Moreover, application of the D_1_R-selective antagonist SCH (10 μM) eliminated the stimulus-evoked response, confirming that the response is due to DA binding to the sensors. Consistent with our results obtained with cultured cells, we found that both the gDA3m and rDA3m sensors had significantly improved sensitivities and responses compared to the corresponding previous-generation sensors (i.e., gDA2m and rDA1m, respectively) (Fig. 3c-e).

**Fig. 3.**
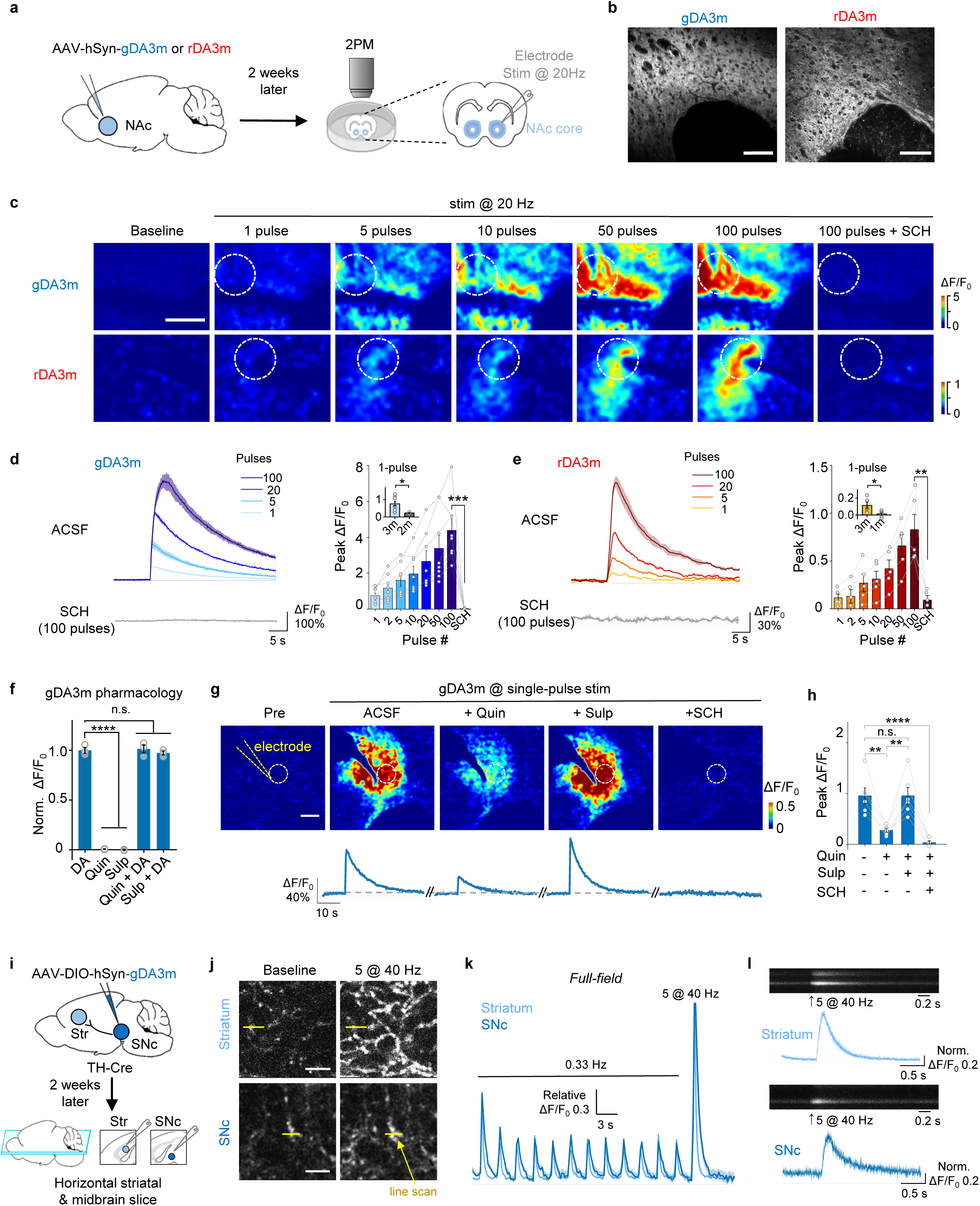
GRAB_DA_ sensors enable endogenous DA detection in mouse acute brain slices a, Schematic illustration depicting the experimental design for panel **b-e**. **b**, Representative fluorescence images showing gDA3m or rDA3m expression in the NAc region. Scale bar, 100 μm. **c**, Example fluorescence response to indicated electrical stimulation measured in the sensor-expressing brain slices. The dashed circles indicate the ROI used to analyze the responses. Scale bar, 100 μm. **d-e**, Representative traces (left) and group summary (right) of the change in sensor fluorescence in response to electrical stimulation (1 ms per pulse). *n*=8 slices from 5 mice (8/5) for gDA3m and *n*=5/4 for rDA3m. The insets show quantification of the ΔF/F_0_ of indicated sensors in response to 1 pulse stimulation. Data replotted from previous results of DA2m and rDA1m^16^. Two-tailed Student’s t-test was performed. p=0.0429 and 0.0138 for gDA3m and rDA3m, p=0.0076 between gDA3m and gDA2m, p=0.0008 between rDA3m and rDA1m. **f**, Normalized fluorescence change in gDA3m to the indicated compounds (each at 1μM). Sulpiride (Sulp), D_2_R antagonist. n=3 wells with 500-600 cells per well. One-way Anova, post hoc Dunnett’s test was performed. p=0.9813, 0.8848 between DA and DA+Quin, or DA+Sulp. **g**. Representative pseudocolored images (top) and traces (bottom) of the fluorescence response of gDA3m to electrical stimuli (5V, 3ms) in ASCF and following drug treatments (each at 1μM). The yellow dashed line indicates the electrode placement. The dashed circles indicate the ROI used to analyze the responses. Scale bar, 100 μm. **h**, Group summary of fluorescence response of gDA3m to electrical stimulation in either ACSF or the indicated drugs. *n*= 6 slices from 5 mice. One-way ANOVA, post hoc Tukey’s test was performed. p=0.00233, p>0.99, p<0.0001 between ACSF and Quin, Sulp or SCH; p=0.00231 between Quin and Sulp. **i**, Schematic illustration depicting the experiment design of panel j-l. **j**, Representative fluorescence images showing gDA3m expression in the striatum (top) and SNc (bottom) in control condition (left) and when delivering 5-pulse stimulation at 40 Hz (right). Scale bar, 5 μm. The yellow lines indicate the hotspot for line-scanning. **k**, Full-frame fluorescence response in response to a series of indicated electrical stimulation (0.5 ms per pulse). The signals were relative to the peak of the first stimulation. Data are shown as mean±SEM. *n* = 6 slices for striatum, *n* = 9 slices for SNc. **l**, Representative line-scan (500 Hz) image in the striatum (top) and the SNc (bottom) and averaged normalized fluorescence traces in response to multi-pulses 40 Hz stimulus. Data are shown as mean±SEM. *n* = 6 areas for striatum and SNc, respectively.

Because the fluorescence of the D_1_R-based gDA3m sensor is not affected by D_2_R-specific compounds such as the D_2_R-specific agonist quinpirole or the D_2_R-specific antagonist sulpiride (Fig. 3f), we examined the effect of D2 autoreceptor activity on DA release in slices expressing the D_1_R-based gDA3m sensor. Activation of endogenous D_2_Rs by the D2-specific agonist quinpirole decreased the stimulus-evoked change in gDA3m fluorescence (Fig. 3g and 3h), reflecting presynaptic inhibition via D2 autoreceptors; this decrease was reversed by the addition of sulpiride, and adding the D_1_R-specific antagonist SCH abolished the stimulation-evoked response (Fig. 3g and 3h).

To measure DA release in both the cell body and terminals of midbrain DA neurons, we injected AAV expressing gDA3m into the substantia nigra pars compacta (SNc), driving expression in both the SNc cell bodies and dopaminergic terminals; we then prepared acute brain slices (Fig. 3i). We found that low-frequency stimulation (0.33 Hz) elicited time-locked transient increases in fluorescence in both the striatum and SNc, while a 40-Hz train of 5 pulses induced a large transient increase in fluorescence, with the signal decay following a slower time course in the SNc compared to the striatum (Fig. 3k). To measure the kinetics of these transients, we performed line-scan microscopy (2 ms/line). Our analysis revealed that the increase in fluorescence upon high-frequency (40 Hz) stimulation had a half-rise time (rise t_1/2_) of 20 ± 10 ms and 22 ± 9 ms in the striatum and SNc, respectively; in contrast, the fluorescence signal decayed to baseline significantly slower in the SNc (decay t_1/2_) compared to the striatum, with decay τ_1/2_ values of 209 ± 49 ms and 125 ± 32 ms, respectively (Fig. 3l). This difference in the time course of DA levels may be attributed—at least in part—to differences in dopamine transporter (DAT) expression between the dorsal striatum and midbrain^35^.

### Validation of our next-generation DA sensors *in vivo*

Next, we examined whether the increased sensitivity of our DA sensors might be suitable for recording *in vivo* DA release in the medial prefrontal cortex (mPFC), which receives relatively sparse dopaminergic innervation from the ventral tegmental area (VTA). We therefore virally expressed the optogenetic tool ChrimsonR (ref. ^36^) in the VTA and either gDA3h or dLight1.3b in the mPFC; we then optically stimulated the VTA and used fiber photometry to measure the signal in the mPFC (Extended Data Fig. 7). We found that activating VTA neurons elicited robust, transient increases in fluorescence in the gDA3h-expressing mPFC, and this increase was blocked by the D_1_R-antagonist SCH (Extended Data Fig. 7b and 7c). In contrast, activating VTA neurons had virtually no effect on dLight1.3b (Extended Data Fig. 7d), indicating that this previous-generation DA sensor lacks the sensitivity to detect *in vivo* DA release in the mPFC. We also found that the gDA3h sensor had discrete, pulse-dependent responses to optogenetic stimulation, and with just one light pulse sufficient to induce a response; in contrast, the less sensitive DA sensor dLight1.3b did not respond in a light pulse number‒dependent manner (Extended Data Fig. 7e-g).

To measure the *in vivo* performance of our new red-shifted GRAB_DA_ sensors, we expressed either rDA3m or rDA3mut in the central amygdala (CeA)—a target of dopaminergic projections from the VTA^37,38^—and the light-activated channel Channelrhodopsin-2 (ChR2, ref. ^39^) in the VTA. We found that activating VTA neurons induced robust, transient increases in rDA3m fluorescence in response to 1-s, 5-s, and 10-s light pulses, with the amplitude of the response increasing incrementally with pulse duration. These responses were virtually eliminated by SCH administration and were absent in mice expressing the DA-insensitive rDA3mut sensor (Extended Data Fig. 8b-d). We also expressed either rDA2m or rDA2mut in both the mPFC and the NAc to measure DA release in these regions in response to VTA activation (Extended Data Fig. 8e); for these experiments, the mice were lightly anesthetized to reduce the tonic activity of dopaminergic neurons. Under these conditions, activating ChR2-expressing VTA neurons reliably induce pulse number‒dependent increases in rDA2m fluorescence in both the densely innervated NAc and the sparsely innervated mPFC, with larger signals induced in the NAc compared to the mPFC; moreover, no signal was detected when we expressed the DA-insensitive rDA2mut sensor (Extended Data Fig. 8f-h). Collectively, these results provide compelling evidence that our next-generation DA sensors can be used *in vivo* to report DA dynamics in several brain regions in real time with high temporal resolution.

To compare the performance of the next-generation DA sensors with the reported GRAB variants, we measured the DA dynamics in the NAc of water-restricted mice when receiving water rewards (Extended Data Fig. 9a and 9b). We found that unpredicted water delivery induced a much larger fluorescence increase of both gDA3m and rDA3m in the NAc compared to gDA2m and rDA1m, respectively (Extended Data Fig. 9c-h). With improved SNR, the gDA3m and rDA3m could readily represent the reward value as the fluorescence response increase with the size of water-drop accordingly. We next compared the performance of rDA3m and RdLight1 by expressing these sensors in opposite sides of the NAc core, and performed bilateral fiber photometry recording (Extended Data Fig. 9i and 9j).

The rDA3m sensor had a substantially higher fluorescence change than RdLight1 across all water-rewarded sessions (Extended Data Fig. 9k and 9l). Taken together, the new DA probes enable DA detection with improved sensitivity and precision *in vivo*.

### Simultaneous *in vivo* imaging of DA and either intracellular cAMP or endocannabinoid signaling during natural behavior

Next, we capitalized on the spectral compatibility between our red-shifted DA sensors and green fluorescent sensors in order to monitor multiple signaling events simultaneously in the same location. The NAc plays a key role in reward processing. Therefore, we measured extracellular DA while also measuring intracellular cyclic AMP (cAMP), the downstream messenger activated by DA receptors and a point of convergence for GPCR signaling^40,41^. We expressed both rDA3m and the green fluorescent cAMP indicator G-Flamp1 (ref. ^42^) in the NAc of male mice and measured both signals during mating, a naturally rewarding condition^43^ (Fig. 4a and 4b). We found that the fluorescence of rDA3m and G-Flamp1 measured in the NAc increased while the male was sniffing the female, mounting the female, during intromission, and during ejaculation (Fig. 4c), with a similar half-rise time of 616 ± 40 ms and 698 ± 53 ms, respectively. Interestingly, however, we found that during all four mating stages the rDA3m signal preceded the G-Flamp1 signal by approximately 200 ms (Fig. 4d and 4f). Moreover, a session-wide cross-correlation analysis revealed that the intracellular cAMP levels measured using G-Flamp1 were closely correlated with the DA signal (Fig. 4e and 4f).

**Fig. 4.**
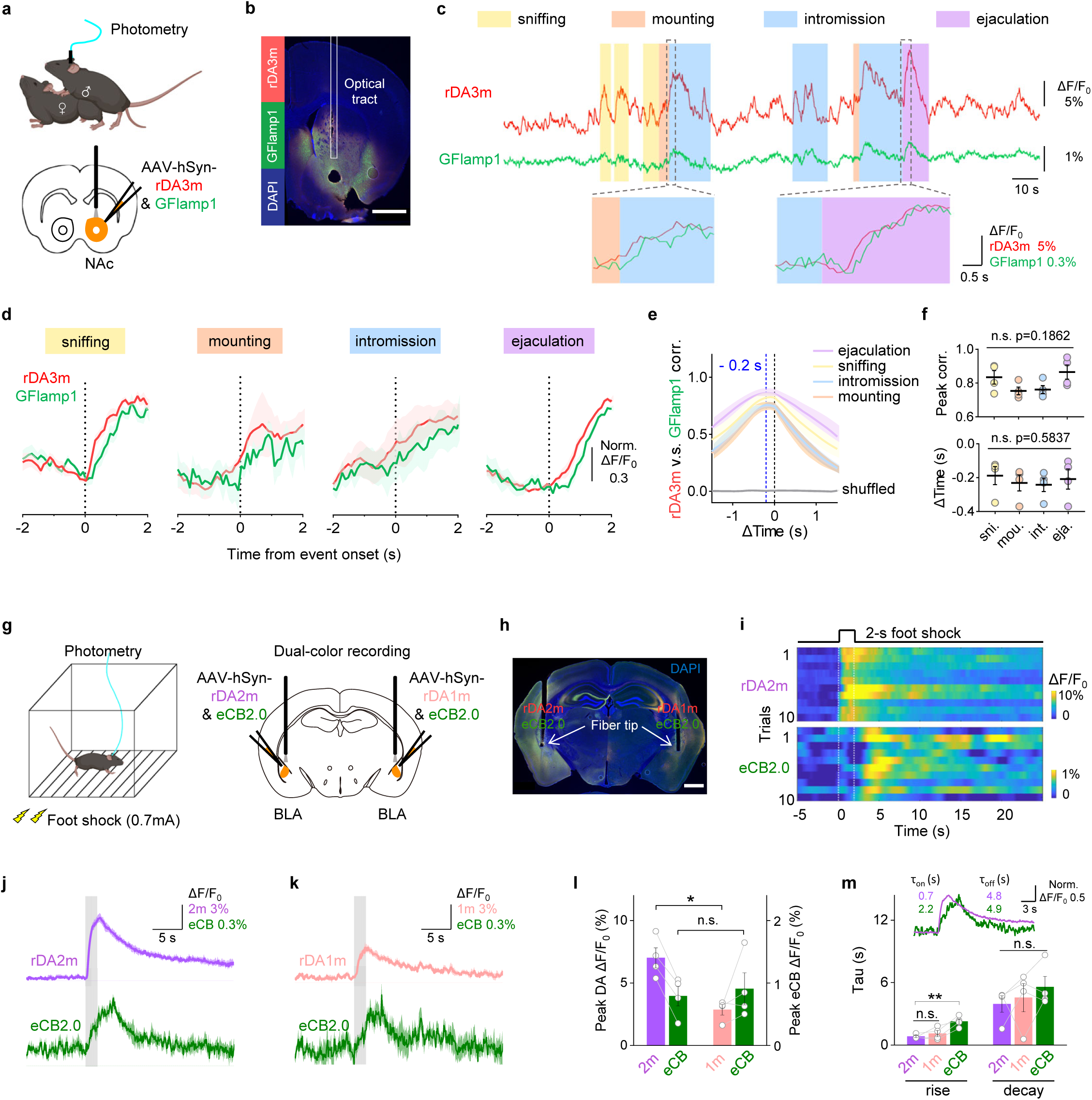
Multiplexed measurements of DA and other neurochemical signals during natural behaviors. **a**, Schematic illustration depicting the experimental design for panel **b-f.** **b**, Histological verification of rDA3m and GFlamp1 expression in NAc. DAPI, 4,6-diamidino-2-phenylindole. Scale bar, 1 mm. **c,** Example traces (top) and zoom-in traces of rDA3m (red) and GFlamp1 (green) signals (*ΔF/F_0_*) simultaneously measured during the indicated stages of mating. **d**, Group-averaged rDA3m and GFlamp1 fluorescence aligned to event onset for all mice. The signals were normalized to respective maxima and minimum. *n*=4 mice. **e**, Cross-correlation between simultaneously recorded rDA3m and GFlamp1 signals during indicated stages and of shuffle group. *n* = 4 mice. **f**, Group summary of peak correlation coefficient (top) and time lag of cross-correlation peak (bottom) between rDA3m and GFlamp-1 signals across mating stages. The black lines indicate mean±SEM. *n* = 4 mice. One-way ANOVA, post hoc Tukey’s test was performed. **g**, Schematic illustration depicting the experimental design for panel **g-k.** **h**, Histological verification of rDA2m and eCB2.0 expression (left side), and rDA1m and eCB2.0 expression (right side) in BLA. Scale bar, 1mm. **i**, Pseudocolored fluorescence responses of rDA2m and eCB2.0 simultaneously measured in the BLA to ten consecutive 2-s foot shock at 0.7 mA. **j**, Average traces of the change in rDA2m (top) and eCB2.0 (bottom) fluorescence from a mouse. The grey shaded area indicates the application of electrical foot shock. Data are shown as mean±SD. **k**, Same as (**h**) with simultaneously recorded contralateral rDA1m and eCB2.0 signals. **l**, Group summary of the peak change in fluorescence of indicated sensors to 2-s foot shock. Peak responses were calculated as the maxima during 0-5 s after foot shock initiation. *n*=4 mice. Paired two-tailed Student’s t-test was performed. *p=0.0390 between rDA2m and rDA1m; p=0.7019 between eCB2.0 groups. **m**, Summary of rise and decay time constants measured for the fluorescence change of indicated sensors in response to foot shock. The inset shows the example average trace of rDA2m and eCB2.0 signals that were normalized to respective maxima and minimum. *n*=4 mice. One-way ANOVA, post hoc Dunnett’s test was performed. Rise time, p=0.5742 and 0.0087 between rDA2m and rDA1m, or eCB2.0. Decay time, p=0.2180 for all groups.

We next examined the extracellular crosstalk between dopaminergic activity and other neurotransmitters. In the brain, the basolateral amygdala (BLA) plays an important role in mediating the fear response and processing aversive memories^44^, and previous studies have shown that both DA signaling and the endocannabinoid (eCB) system in the BLA participate in anxiety and fear formation^45^. However, the relative timing of DA and eCB signals under stress conditions remains unknown, particularly at high temporal resolution. We therefore expressed both rDA2m and the green fluorescent eCB indicator eCB2.0 (ref. ^46^) in the BLA in one hemisphere and measured both signals while applying mild foot shocks to induce stress; as a control, we also expressed and measured rDA1m and eCB2.0 in the other hemisphere (Fig. 4g and 4h). We found that rDA2m and eCB2.0 had reproducible, time-locked transient increases in fluorescence upon delivery of a 2-s foot shock; moreover, although the signal produced by eCB2.0 was similar between hemispheres, the signal produced by rDA2m was approximately twice as large as the signal produced by rDA1m (Fig. 4i-l). We also examined the kinetics of the DA and eCB signals and found that although the τ_off_ rates were similar for DA and eCB (on a order of 4-5 s), the τ_on_ rate was significantly faster for DA (∼0.8 s) compared to eCB (∼2.2 s), with no significant difference between rDA2m and rDA1m (Fig. 4m). This difference between the relatively rapid DA signal and the slower eCB signal is in consistent with the known signaling mechanism of small-molecule transmitters such as DA and lipid neurotransmitters such as eCBs^47,48^.

### Simultaneously measuring DA and ACh release *in vivo* during an auditory Pavlovian conditioning task

Dopaminergic signaling plays a key role in reinforcing learning and memory through the mesocorticolimbic system^49–51^. External rewards such as food also elicit characteristic changes in acetylcholine (ACh) levels that promote learning and motivate action^52^. However, the relationship between DA release and ACh release—as well as the dynamics of their release in the mesocorticolimbic system during reinforcement learning—are poorly understood. We therefore measured the release of both DA and ACh during auditory Pavlovian conditioning tasks by co-expressing rDA3m and the green fluorescent ACh sensor ACh3.0 (ref. ^53^) in both the NAc and the mPFC. Mice were head-fixed, water-restricted, and trained to associate a specific auditory cue with either a water reward (associated with tone A) or a punitive mild puff of air applied to the eye (associated with tone B) (Fig. 5a and 5b). Initially, rDA3m mainly responded to the reward, while ACh3.0 responded both to the reward and the punishment, with minimal response to the auditory cues (Fig. 5c). After five days of training, however, the mice selectively associated the stimulus-predicted cue (tone A or tone B) with the subsequent delivery of reward or punishment (i.e., water or air puff, respectively). The rDA3m and ACh3.0 signals in the NAc of these trained mice increased in response to the water-predicted tone, but decreased in response to the punishment-related tone, whereas their responses to actual reward or punishment were remained in the current paradigm (Fig. 5c and 5e-f). The development of excitatory responses to the reward cue and inhibitory responses to the punishment cue is consistent with the so-called reward-prediction-error theory^51^. Interestingly, rDA3m and ACh3.0 signals in the mPFC increased in response to both stimulus-predictive cues and the actual outcomes of both valences in naive mice. After training sessions, unlike what is usually seen in reward prediction error patterns, there was no signal shift (Fig. 5d-f). Furthermore, within brain areas (NAc or mPFC), DA and ACh signals were positively correlated with each other during reward and punishment trials (Fig. 5c and 5d), indicating that a similar upstream process regulates DA and ACh release in these two brain regions or a local neuromodulatory effect that one enhances the other^20^. However, these signals were not correlated between brain areas (NAc and mPFC; Fig. 5g), suggesting a heterogeneity of neurotransmission in the mesocorticolimbic system. As a control, we found that systemic administration of the D_1_R blocker SCH significantly reduced the rDA3m signal but did not affect the ACh3.0 signal (Extended Data Fig. 10).

**Fig. 5.**
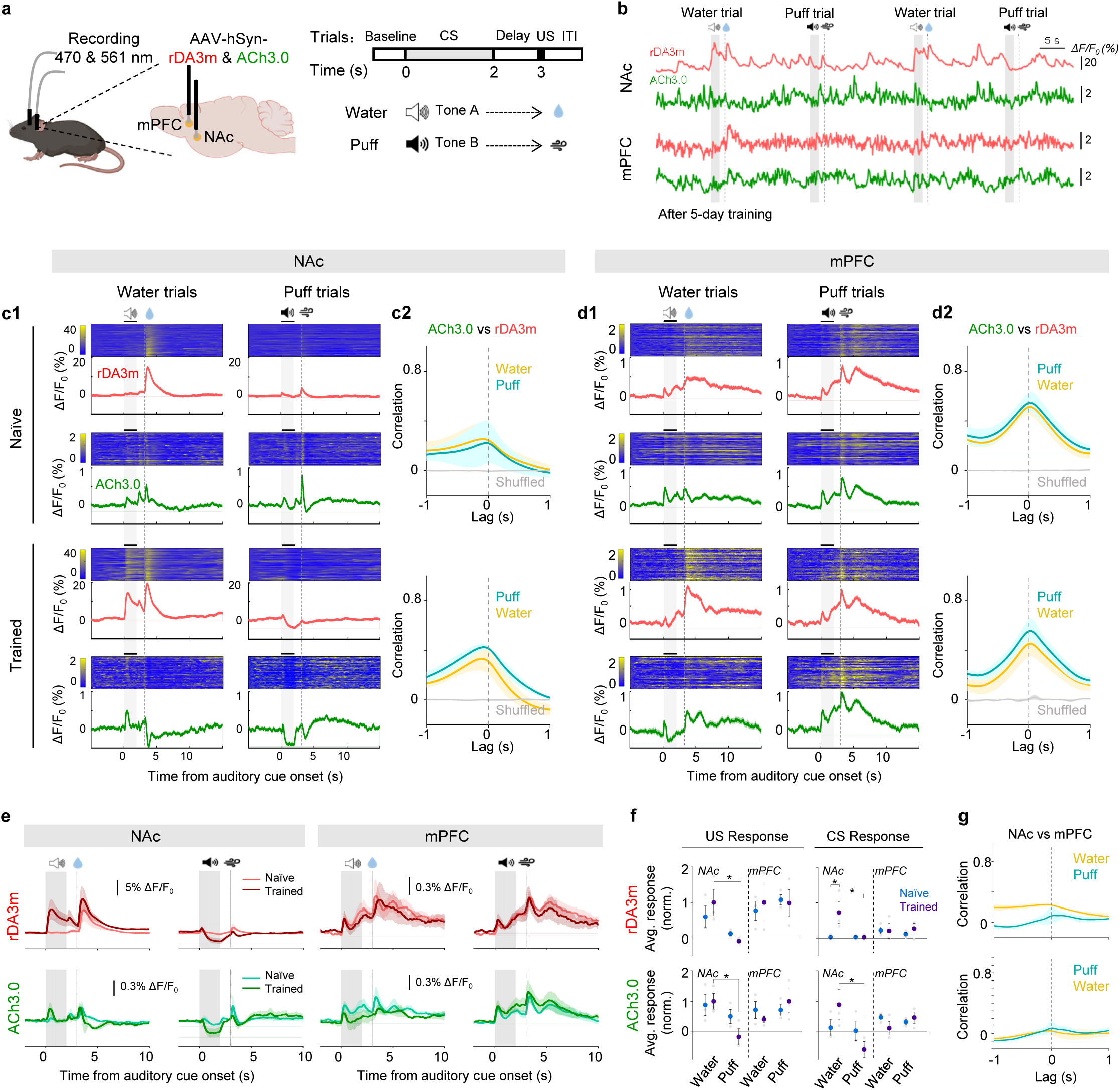
DA and ACh signals in mouse NAc and mPFC during an auditory Pavlovian conditioning task. **a**, Schematic illustration depicting experimental design for panel **b-g**. **b**, Example traces of rDA3m (red) and ACh3.0 (green) signals (ΔF/F_0_) simultaneously measured in the NAc (top) and mPFC (bottom) from a trained mouse during four consecutive trials. The audio, water and puff delivery are indicated above. **c**, Representative time-aligned pseudocolored images and averaged traces of rDA3m and ACh3.0 fluorescence from a mouse in naïve (top) and trained (bottom) state (**c1**). Shown are 100 consecutive trials (mean±SD) in one mouse. The grey shaded area indicates the application of audio. The dashed line indicates the delivery of water or puff. Session-wide correlation between rDA3m and ACh3.0 signals across naïve (top) and trained sessions (bottom) **(c2)**. *n*=3 mice. **d**, same as (**c**) with simultaneously recorded rDA3m and ACh3.0 signals in the mPFC. **e**, Group-averaged rDA3m (top) and ACh3.0 (bottom) fluorescence in the NAc (left) and mPFC (right) for all mice under naïve and trained state. Water or puff sessions are indicated above. *n*=3 mice. **f**, Group analysis of the normalized average change of rDA3m (top) and ACh3.0 (bottom) signals to US (left) and CS (right) in different sessions. The average response was calculated as the average ΔF/F_0_ in the 1 s after the behavior onset. The grey points indicate data from individual animal; Average and SEM are shown by data points with state-represented color. *n*=3 mice. Two-way ANOVA, post hoc Sidak’s test was performed between water and puff sessions and between naïve and trained state. Responses in the trained mice NAc, p=0.0145 (rDA3m-US), 0.0287 (rDA3m-CS), 0.0371 (ACh3.0-US), 0.0356 (ACh3.0-CS) between water and puff trial; rDA3m water trial CS response in the NAc, p=0.0290 between naïve and trained. **g**, Session-wide cross-correlation between rDA3m (top) or ACh (bottom) signals recorded in the NAc and mPFC. *n*=3 mice.

### Spatially resolved imaging of cortical DA release

Dopaminergic signaling also plays a key role in modulating several physiological processes, including motor control and reward perception. The cortex receives dopaminergic innervation from both the SNc and VTA, which send distinct dopaminergic signals^7,54–56^. To test whether our high-affinity gDA3h sensor can be used to monitor behavior-related changes in cortical DA levels with high spatiotemporal resolution, we expressed gDA3h in the M1/M2 motor cortex (Fig. 6a) and performed head-fixed *in vivo* two-photon imaging (Fig. 6b). As DA is thought to be a key regulator of locomotion and aversive events^4,57,58^, during imaging, the mouse was placed on a treadmill and gDA3h fluorescence was measured in response to a 70-s bout of forced running (Fig. 6c), an electrical tail shock (Fig. 6d), or an auditory stimulus (Fig. 6e). Interestingly, we observed a robust, rapid, reproducible increase in gDA3h fluorescence aligned to the onset of forced running and tail shock, but not in response to the auditory stimulus (Fig. 6c-g). Similar results were obtained when we expressed gDA3m, whereas dLight1.3b was not sufficiently sensitive to capture these relatively mild changes in DA (Extended Data Fig. 11). As a negative control, no response was measured in mice expressing membrane-targeted EGFP (Fig. 6c and 6g; Extended Data Fig. 10). We then examined the spatial patterns of DA release during forced running and foot shock on a trial-by-trial basis; interestingly, using select regions of interest (ROIs) we observed distinct patterns during running and shock (Fig. 6c-e). Consistent with this observation, we identified four distinct categories of cell-sized ROIs by performing hierarchical cluster analysis to analyze the average response of individual ROIs (Fig. 6h-j). All four categories were observed in all animals tested (Fig. 6k-m). Although most areas had no response, a small subset of responsive regions (representing 0.7% of the entire area) had increases in DA levels during both running and shock, while 3.61% and 3.68% of the entire area were associated exclusively with running or shock, respectively (Fig. 6n-o). Taken together, these results show that our gDA3h sensor can be used to map spatially and functionally heterogeneous patterns of DA release in the motor cortex at subsecond resolution.

**Fig. 6.**
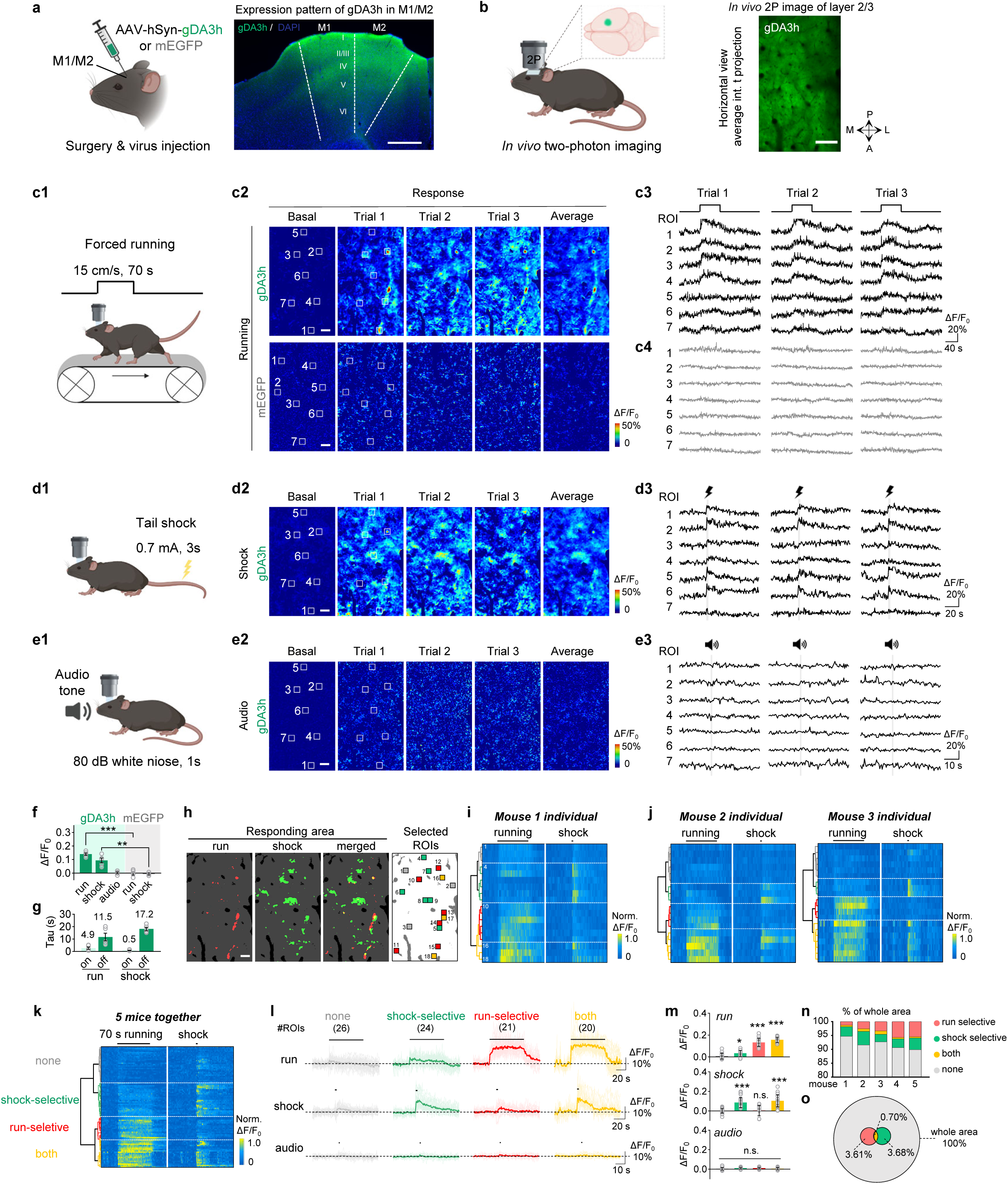
Spatially resolved heterogeneous cortical DA dynamics in mice. **a-b**, Schematic illustration depicting the strategy for virus injection and head-fix two-photon imaging in the mouse motor cortex. Example fluorescence image showing gDA3h expression in the M1/M2 region in a coronal brain slice (**a**). Scale bar, 500 μm. Representative in vivo 2P image of the layer 2/3 in the M1/M2 cortex showing gDA3h fluorescence (**b**). Scale bar, 100 μm. **c**, Schematic cartoon illustrating the forced running experiments (**c1**), representative pseudocolored response images (**c2**) and traces measured at indicated ROIs during three consecutive trials (**c3**) in the head-fix mice expressing gDA3h (top) or membrane-targeted EGFP (mEGFP, bottom). The white squares indicate ROI to analyze signals. Scale bar, 50 μm. **d-e**, Similar to (**c**) except mice were subjected to 3-s tail shock, 0.7mA (**d**) or 1-s audio stimulation (**e**). Two-photon imaging was performed in the same region across different behaviors. **f**, Group summary of the peak fluorescence change measured in the motor cortex in mice expressing gDA3h or mEGFP in response to indicated stimulus. *n*=5 mice for gDA3h and 4 mice for mEGFP. Two-tailed Student’s t-test was performed. p<0.0001 between gDA3h and mEGFP for running and p=0.0017 for shock. **g**, Group summary of the rise and decay time constant of the gDA3h signals in response to forced running and tail shock. *n*= 5 mice. **h**, Example image showing the spatial responding pattern to forced running, tail shock, merge and ROI selection. The responding area was defined according to fluorescence signals during indicated behaviors. ROIs (30 μm x 30 μm) were randomly selected inside and outside of responding area and colored according to the type of fluorescence responses. **i**, Hierarchical clustering of ROI-specific responses to running and shock from a mouse. ROIs were indicated in (**h**). **j**, Same to (**h**) with data collected from another two mice. **k**, Population data showing hierarchical clustering of ROI-specific response. *n*=91 ROIs from 5 mice. **l-m**, Average (bold lines) and individual (thin lines) traces (**l**) and quantifications of response amplitudes (**m**) from ROIs within each cluster in (**k**) during different behaviors. The black lines indicate the application of indicated stimulus. *n*=26, 24, 21 and 20 ROIs from 5 mice for each cluster. One-way ANOVA, post hoc Tukey’s test was performed. Forced running, p=0.0128 between *none* and *shock*; p=0.0585 between *running* and *both*. Tail shock, p=0.9989 between *none* and *running*; p=0.6474 between *shock* and *both*. Audio, n.s. p=0.0766 for all clusters. **n**, Percentage of area that was responsive to the indicated stimulus as in (**h**). **o**, Venn diagram of the imaged motor cortex area that was responsive to the indicated stimulus. Data collected from 5 mice.

## DISCUSSION

Here we used rational design to develop a third-generation series of highly sensitive and highly selective DA sensors suitable for use in *in vivo* multiplex imaging. Our improved red fluorescent DA sensors performed as well as their corresponding green fluorescent counterparts in terms of their sensitivity at detecting DA signals, thereby narrowing the performance gap between red and green fluorescent DA sensors. Moreover, the features of these optimized sensors are compatible with other recently developed optical sensors (e.g., cAMP, eCB, and ACh sensors) for use in monitoring signaling events in the central nervous system in real time.

Although several fluorescent DA sensors have been developed using the GPCR-based strategy, the sensors reported to date lack the sensitivity needed to monitor DA release in brain regions with relatively sparse dopaminergic innervation or individual release events, yielding only to the small changes in fluorescence; our highly sensitive series of new DA sensors overcomes this limitation. Moreover, our new series of GRAB_DA_ sensors can be used to monitor DA dynamics *in vivo* in several brain regions such as the NAc, amygdala, and cortex. Importantly, our simultaneous recordings of localized DA release in the NAc and mPFC revealed a lack of synchronized DA release from distinct axonal termini. Specifically, in the context of Pavlovian conditioning the well-known theory of prediction-error was observed in the NAc, but not in the mPFC; in the mPFC, the DA signals were consistent with the reported cortical dopaminergic activity in the context of stimulus discrimination^59^. This difference in DA dynamics between these two brain regions suggests functional heterogeneity within VTA dopamine neurons, as has been indicated by unique intrinsic properties of cortex-projecting dopamine neurons^60^. In addition, even in a given brain region such as the motor cortex, our two-photon imaging of gDA3h revealed behaviorally related, spatially resolved heterogeneity in cortical dopaminergic signaling.

Our medium-affinity DA sensors (gDA3m, rDA2m, and rDA3m) are particularly well suited for imaging DA dynamics in brain regions that contain moderate or high DA levels and for monitoring rapidly changing events that require a rapid off rate. In contrast, our high-affinity sensors (gDA3h, rDA2h, and rDA3h) can be used to monitor small changes in DA levels, for example in sparsely innervated brain regions. The improved signal-to-noise ratio and sensitivity of these sensors have the potential to facilitate the detection of individual release events, thereby greatly enhancing our understanding of the biophysical characteristics of DA release. In addition, the distinct pharmacological profiles of these receptor subtype‒based sensors allow for DA imaging while manipulating the activity of specific receptors. These properties are also valuable for studying DA pharmaceutical agents and for screening compounds that target specific DA receptor subtypes.

In combination with multicolor fluorescence imaging of other signaling events, our new series of DA sensors can be used to functionally map neurochemical activity. Moreover, this robust set of GRAB_DA_ sensors will help pave the way to a deeper understanding of the complexity of the dopaminergic system.

## METHODS

### Animals

All procedures for animal surgery and experimentation were performed in accordance and approved by the laboratory animal care and use committees of Peking University, the Institutional Animal Care and Use Committee at Oregon Health and Science University, the National Institutes of Health Guide for the Care and Use of Laboratory Animals and Harvard Animal Care and Use Committee. Both male and female postnatal day 0 (P0) Sprague-Dawley rats were used to prepare cultured cortical neurons; P48-P90 wild-type C57BL/6N mice (Beijing Vital River Laboratory), wild-type C57BL/6J mice (Beijing Vital River Laboratory), TH-Cre mice (The Jackson Laboratory; B6. Cg-*7630403G23Rik^Tg(Th-cre)1Tmd^*/J) and DAT-Cre mice (The Jackson Laboratory; B6.SJL-Slc6a3^tm1.1(cre)Bkmn^/J) were used in this study. All animals were housed at 18–23°C in 40–60% humidity under a normal 12-h light–dark cycle with food and water available *ad libitum*.

### AAV expression

AAV2/9-hSyn-gDA3m (2.8 × 10^13^ viral genomes (vg) per ml), AAV2/9-hSyn-gDA3h (8.43 × 10^13^ vg/ml), AAV2/9-hSyn-gDA3mut (6.21 × 10^13^ vg/ml), AAV2/9-hSyn-gDA2m (5.72 × 10^13^ vg/ml), AAV2/9-hSyn-dLight1.3b (6.49 × 10^13^ vg/ml), AAV2/9-DIO-hSyn-gDA3m (6.2 × 10^13^ vg/ml), AAV2/9-EGFP-CAAX(3.5 × 10^13^ vg/ml), AAV2/9-hSyn-ACh3.0 (8.0 × 10^13^ vg/ml), and AAV2/9-hSyn-eCB2.0 (9.2 × 10^13^ vg/ml), AAV2/9-hSyn-GFlamp1 (7.29 × 10^13^ vg/ml) were packaged at Vigene Biosciences. AAV2/9-hSyn-rDA1m (1.04 × 10^13^ vg/ml), AAV2/9-hSyn-rDA2m (6.04 × 10^12^ vg/ml), AAV2/9-hSyn-rDA2h (5.31 × 10^12^ vg/ml), AAV2/9-hSyn-rDA2mut (5.09 × 10^12^ vg/ml), AAV2/9-hSyn-rDA3m (3.29 × 10^12^ vg/ml), AAV2/9-hSyn-rDA3h (6.36 × 10^12^ vg/ml), AAV2/9-hSyn-rDA3mut (6.16 × 10^12^ vg/ml), AAV2/9-hSyn-RdLight1 (6.12 × 10^12^ vg/ml), AAV2/9-hSyn-hChR2(H134R)-eYFP (5.49 × 10^12^ vg/ml), and AAV2/9-hSyn-ChrimsonR-tdTomato (2.52 × 10^12^ vg/ml) were packaged at BrainVTA). Where indicated, the AAVs were either used to infect cultured neurons or were injected *in vivo* into specific brain regions.

### Molecular biology

cDNAs encoding the various DA receptors were cloned from the respective human (hORFeome database 8.1), bovine, sheep, waterbear, bat, cat, monkey, zebrafinch and red fire ant genes (Shanghai Generay Biotech). DNA fragments were PCR-amplified using specific primers (Tsingke Biological Technology) with 25-30-bp overlap. Plasmids were generated using Gibson assembly^61^, and all plasmid sequences were verified using Sanger sequencing. For screening and characterization in HEK293T cells, the green and red fluorescent DA sensors were cloned into the pDisplay vector (Invitrogen). The IRES-mCherry-CAAX cassette (for expressing green fluorescent sensors) or IRES-EGFP-CAAX cassette (for expressing red fluorescent sensors) was inserted downstream of the sensor gene to serve as a cell membrane marker and to calibrate the sensor’s fluorescence. Site-directed mutagenesis was performed using primers containing randomized NNB codons (48-51 codons in total, encoding all 20 amino acids; Tsingke Biological Technology) or defined codons. For characterization in cultured neurons, the sensor gene was cloned into a pAAV vector under the control of the human synapsin (*SYN1*) promoter (*pAAV-hSyn*). For luciferase complementation assay, the receptor-SmBit or sensor-SmBit was generated from β_2_AR-SmBit^33^. For the Tango assay, genes encoding the wild-type receptors or the indicated sensors were cloned into the pTango vector^34^.

### Cell culture

HEK293T cells were cultured at 37°C in humidified air containing 5% CO_2_ in DMEM (Biological Industries) supplemented with 10% (vol/vol) FBS (Gibco) and 1% penicillin-streptomycin (Gibco). For experiments, the cells were seeded in 96-well plates or on 12-mm glass coverslips in 24-well plates. At 60-70% confluency, the cells were transfected with a mixture of polyethylenimine (PEI) and plasmid DNA at a 3:1 (w/w) ratio; the culture medium was replaced with fresh medium 6-8 h after transfection, and imaging was performed 24-48 h after transfection. Rat cortical neurons were prepared from P0 Sprague-Dawley rats. In brief, cortical neurons were dissociated from the dissected rat cerebral cortex by digestion in 0.25% trypsin-EDTA (Biological Industries) and then plated on poly-D-lysine-coated (Sigma-Aldrich) 12-mm glass coverslips in 24-well plates. The neurons were cultured in Neurobasal medium (Gibco) containing 2% B-27 supplement (Gibco), 1% GlutaMAX (Gibco), and 1% penicillin-streptomycin (Gibco) at 37°C in humidified air containing 5% CO_2_. The cultured neurons were transfected with AAVs expressing the indicated sensors at 3-5 days *in vitro* (DIV3-5) and imaged at DIV11-14.

### Fluorescence imaging of cultured cells

Before imaging, the culture medium was replaced with Tyrode’s solutions containing (in mM): 150 NaCl, 4 KCl, 2MgCl_2_, 10 HEPES, and 10 glucose (pH adjusted to 7.35-7.45 with NaOH). The cells then were imaged in a custom-made chamber using an inverted Ti-E A1 confocal microscope (Nikon) and an Opera Phenix high-content screening system (PerkinElmer). The confocal microscope was equipped with a 10x/0.45 number aperture (NA) objective, a 20x/0.75 NA objective, a 40x/1.35 NA oil-immersion objective, a 488-nm laser, and a 561-nm laser. Green fluorescence was collected using a 525/50-nm filter, and red fluorescence was collected using a 595-50nm filter. During imaging, the following compounds were applied via bath application or via a custom-made perfusion system at the indicated concentrations: DA (Sigma-Aldrich), SCH (Tocris), Etic (Tocris), SKF (Tocris), Quin (Tocris), Glu (Sigma-Aldrich), GABA (Tocris), L-Dopa (Abcam), ACh (Solarbio), 5-HT (Tocris), HA (Tocris), OA (Tocris), NE (Tocris), and Sulp (MedChemExpress). To measure the kinetics of the GRAB_DA_ sensors, the confocal line-scanning mode (2,600 Hz) was used to record the fluorescence response when the cells were locally puffed with DA via a pipette positioned at the cells. Similarly, the decay kinetics were measured by locally puffing cells with the respective antagonist in the presence of saturating DA concentration. The Opera Phenix system was equipped with a 20x/1.0 NA, a 40x/0.6 NA objective, a 40x/1.15 NA water-immersion objective, a 488-nm laser, and a 561-nm laser. Green fluorescence was collected using a 525/50-nm emission filter, and red fluorescence was collected using a 600/30-nm emission filter, and the fluorescence intensity of the red and green fluorescent sensors was calibrated using EGFP and mCherry, respectively.

### Spectra and photoactivation measurements

Plasmids expressing the GRAB_DA_ sensors were transfected into HEK293T cells in six-well plates (for 1-photon spectra) or on 12-mm coverslips (for 2-photon spectra). For measuring the 1P spectra, the cells were harvested 24-30 h after transfection and transferred to 384-well plates in the absence or presence of 100 μM DA. The excitation and emission spectra were then measured at 5-nm increments using a Safire2 multi-mode plate reader (Tecan). The fluorescence measured in non-transfected cells was subtracted as background. The 2P spectra of gDA3m were measured at 10-nm increments ranging from 700-1050 nm using an Ultima Investigator 2-photon microscope (Bruker) equipped with a 20x/1.00 NA water-immersion objective (Olympus) and an InSight X3 tunable laser (Spectra-Physics). The 2P spectra of rDA3m were measured at 10-nm increments ranging from 820-1300 nm using an A1R MP+ multiphoton microscope (Nikon) equipped with a 25×/1.10 NA objective (Nikon) and a Chameleon Discovery tunable laser (Coherent). Laser power was calibrated according to the output power of the tunable 2P laser with various wavelengths. Bursts of 488-nm laser light (1 s duration, 210 μW, ∼0.4 W cm^−2^ intensity) were applied to induce blue light‒mediated photoactivation.

### Luciferase complementation assay

At 50-60% confluency, HEK293T cells were co-transfected with the indicated wild-type receptor or sensor together with the respective LgBit-mG construct. Approximately 24-36 h after transfection, the cells were dissociated using a cell scraper, resuspended in PBS, and transferred to 96-well plates. DA at concentrations ranging from 0.01 nM to 100 μM and 5 μM furimazine (NanoLuc Luciferase Assay, Promega) were then bath-applied to the cells. After a 10-min reaction in the dark at room temperature, luminescence was measured using a VICTOR X5 multi-label plate reader (PerkinElmer).

### Tango assay

The Tango assay was perform as previously described^34^ using HTLA cells (a gift from Bryan L. Roth) seeded in 6-well plates and transfected with plasmids expressing the indicated receptors or sensors. Twenty-four hours after transfection, the cells were collected using trypsin digestion, plated in 96-well plates, and DA was added to the media at concentrations ranging from 0.01 nM to 10 μM. The cells were then cultured for an additional 12 h for luciferase expression; 5 μM Bright-Glo (Fluc Luciferase Assay System, Promega) was then added to the wells, and luminescence was measured using a VICTOR X5 multi-label plate reader (PerkinElmer).

### Two-photon imaging in the NAc in acute mouse brain slices

Adult (6-8 weeks of age) C57BL/6N of both sexes were anesthetized with an intraperitoneal injection of 2,2,2-tribromoethanol (Avertin, 500 mg/kg body weight; Sigma-Aldrich), and AAVs were injected (300 nL per injection site at a rate of 40 nl/min) into the NAc using the following coordinates: AP: +1.4 mm relative to Bregma; ML: ±1.2 mm relative to Bregma; and DV: −4.0 mm from the dura. Two weeks after virus injection, the mice were deeply anesthetized, and the heart was perfused with slicing buffer containing (in mM): 110 choline chloride, 2.5 KCl, 1.25 NaH_2_PO_4_, 25 NaHCO_3_, 7 MgCl_2_, 25 glucose, and 0.5 CaCl_2_. The mice were then decapitated and the brains were immediately removed and placed in cold oxygenated slicing buffer. The brains were sectioned into 300-μm-thick coronal slices using a VT1200 vibratome (Leica), and the slices were incubated at 34°C for at least 40 min in oxygenated artificial cerebrospinal fluid (ACSF) containing (in mM): 125 NaCl, 2.5 KCl, 1 NaH_2_PO_4_, 25 NaHCO_3_, 1.3 MgCl_2_, 25 glucose, and 2 CaCl_2_. Two-photon imaging was performed using either an FV1000MPE 2P microscope (Olympus) equipped with a 25×/1.05 NA water-immersion objective and a mode-locked Mai Tai Ti:Sapphire laser (Spectra-Physics) or an Ultima Investigator 2P microscope (Bruker) equipped with a 20x/1.00 NA objective (Olympus) and an InSight X3 tunable laser (Spectra-Physics). A 920-nm laser was used to excite the gDA3m sensor, and fluorescence was collected using a 495-540-nm filter (for the FV1000MPE microscope) or 490-560-nm filter (for the Ultima Investigator microscope); a 950-nm laser was used to excite the rDA3m sensor, and fluorescence was collected using a 575-630-nm filter (for the FV1000MPE microscope) or a 570-620-nm filter (for the Ultima Investigator microscope). For electrical stimulation, a bipolar electrode (model WE30031.0A3, MicroProbes) was positioned near the NAc core under fluorescence guidance, and imaging and stimulation were synchronized using an Arduino board with custom-written software. The stimulation voltage was set at 4-6 V. Where indicated, compounds were added by perfusion at a flow rate of 4 ml/min.

### Two-photon imaging in the striatum and SNc in acute mouse brain slices

Adult TH-Cre and wild-type mice of both sexes were anesthetized with isoflurane, and the indicated AAVs were injected into the SNc region at the following coordinates: AP: −2.3 mm relative to Bregma; ML: ±1.3 mm relative to Bregma; and DV: −4.5 mm from the dura. After 2-3 weeks, the mice were anesthetized with isoflurane, decapitated, and the brain was removed and placed in warm (32-35°C) extracellular solution containing (in mM): 126 NaCl, 2.5 KCl, 1.2 MgCl_2_, 2.4 CaCl_2_, 1.4 NaH_2_PO_4_, 25 NaHCO_3_, and 11 dextrose; the solution also contained MK-801 (10 µM) to prevent NMDA-mediated excitotoxic damage. Horizontal slices (222-µm thickness) containing the midbrain and striatum were cut using a vibratome in warm extracellular solution and recovered at 30°C for ≥30 min before experiments. Two-photon imaging of the striatal and SNc slices was performed using a custom-built 2-photon microscope with ScanImage software^62^. Full-frame images (128×128 pixels) were captured at a rate of 4 Hz. Line scans through areas of interest were taken at 2 ms/line. Images were analyzed using ImageJ (National Institutes of Health) and custom software written using MATLAB.

### *In vivo* fiber photometry recording in mice

Adult mice were anesthetized with isoflurane, and the indicated AAVs were injected (300 nl total volume) were injected as follows. The NAc was targeted using the following coordinates: AP: +1.4 mm relative to Bregma; ML: ±1.2 mm relative to Bregma; and DV: −4.0 mm from the dura. The mPFC was targeted using the following coordinates: AP: +1.98 mm relative to Bregma; ML: ±0.3 mm relative to Bregma; and DV: −1.8 from the dura. Finally, the BLA was targeted using the following coordinates: AP: −1.4mm relative to Bregma; ML: ±3 mm relative to Bregma; and DV: −4.5 mm from the dura. Optical fibers (200 µm diameter, 0.37 NA; Inper) were implanted using the AAV injection site and secured with resin cement (3M). Two weeks after injections, photometry and animal behaviors were recorded using an FPS-410/470/561 photometry system (Inper). In brief, a 10-Hz (with 20-ms pulse duration) 470/5-nm filtered light-emitting diode (LED) at 20-30 μW was used to excite the green fluorescent sensors, and a 10-Hz (20-ms pulse duration) 561/5-nm filtered LED at 20-30 μW was used to excite the red fluorescent sensors. Alternating excitation wavelengths were delivered, and the fluorescence signals were collected using a CMOS camera during dual-color imaging. To minimize autofluorescence of the optical fiber, the recording fiber was photobleached using a high-power LED before recording. The photometry data were analyzed using a custom-written MATLAB (MATLAB R2022a, MathWorks) program, and background autofluorescence was subtracted from the recorded signals.

#### Unpredicted water reward

Adult female (8-9 weeks in age) C57BL/6J mice were prepared for this experiment. AAV-hsyn-rDA3m or AAV-hsyn-RdLight1 virus (300nl for each virus) was bilaterally injected into the NAc. Intraoral cheek fistula implanted and water-restricted mice freely received water delivery (around 10 ul per trial; 25 trials per session; inter-reward-interval = 20 s). To calculate ΔF/F_0_, baseline was chosen as the average fluorescence signal during 4.5-5.0 s ahead of water delivery.

#### Mating behaviors

Adult (8-9 weeks of age) male C57BL/6N mice were used for these experiments. A mixture of AAV-hsyn-rDA3m (300 nl) and AAV-hsyn-Gflamp1 (300 nl) was injected into the NAc as described above. Experienced adult (8-9 weeks in age) ovariectomized (OVX) female C57BL/6N mice were also used to measure the male’s mating behaviors. Three days before recording, the OVX female mice received intraperitoneal injections of estrogen (50 μl, 0.2 mg/ml on day 1 and 50 μl, 0.1 mg/ml on day 2) or progesterone (50 μl, 1 mg/ml on day 3). The various sexual behaviors were defined as previously described^16^. To calculate *ΔF/F_0_*, the baseline was defined as the average fluorescence measured 1-5 min before introducing the female.

*Foot shock.* Adult (8-9 weeks of age) male C57BL/6N mice were used for these experiments. A mixture of AAV-hsyn-eCB2.0 (300 nl) and either AAV-hsyn-rDA2m or AAV-hsyn-rDA1m (300 nl each) was bilaterally injected into the BLA as described above. The mouse was placed in a shock box, and 5 2-s pulses of electricity at an intensity of 0.7 mA were delivered with an interval of 90-120 s between trials. To calculate *ΔF/F_0_*, the baseline values was defined as the average fluorescence measured during a 2-s window prior to the first shock trial.

#### Pavlovian auditory conditioning task

Adult (8-9 weeks of age) female C57BL/6J mice were used for these experiments. A mixture of AAV-hsyn-gACh3.0 (300 nl) and AAV-hsyn-rDA3m (300 nl) was injected into the NAc and mPFC as described above. A stainless-steel head holder was attached to the skull using resin cement in order to restrain head-fix the animal. For water delivery, an intraoral cheek fistula was implanted in each mouse as previously described^14^. In brief, incisions were made in the cheek and the scalp at the back of the neck. A short, soft silastic tube (inner diameter: 0.3 mm, outer diameter: 0.46 mm) connected via an L-shaped stainless-steel tube was then inserted into the cheek incision site. The steel tube was inserted into the scalp incision, and the opposite end was inserted into the oral cavity. The head-fixed mice were habituated to the treadmill apparatus for 2 days (1 h per day) before the experiments to minimize potential stress. On the day of the experiment, the Pavlovian auditory conditioning task was performed using two pairs of auditory cues and outcomes, with tone A (2.5k Hz, 70 dB, 2 s duration) paired with delivery of 10 μl of 5% sucrose water and tone B (15k Hz, 70 dB, 2 sec duration) paired with deliver of an air puff to the eye. These two pairs were randomly delivered with a 15-20 s randomized inter-trial interval. The water and air puff delivery were precision-controlled using a stepper motor pump and a solenoid valve, respectively. A custom Arduino code was used to control the timing of the pump and solenoid valve and to synchronize the training devices with the photometry recording system. To calculate *ΔF/F_0_*, the baseline was defined as the average fluorescence signals measured 4.5-5.0 s prior to the first auditory cue.

### Fiber photometry recording of optogenetically induced DA release in mice

Adult (8-9 weeks of age) male C57BL/6N mice were used for these experiments. The mice were anesthetized with isoflurane, and AAV-hsyn-gDA3h (300 nl) or AAV-hsyn-dLight1.3b (300 nl) was injected into the mPFC as described above. AAV-hsyn-ChrimsonR-tdTomato (300 nl) was also injected into the VTA using the following coordinates: AP: −2.9 mm relative to Bregma; ML: ±0.65 mm relative to Bregma; and DV: −4.1 mm from the dura. Optical fibers (200-µm diameter, 0.37 NA; Inper) were implanted in the same injection sites and secured with resin cement (3M). Two weeks after virus injection, photometry recording was performed using a commercially available photometry system (Thinker Tech, Nanjing, China).

A 470/25-nm bandpass-filtered (model 65-144; Edmund Optics) LED light (Cree LED) was used to excite the green fluorescent sensors. The emitted fluorescence was bandpass filtered (525/25 nm, model 86-354; Edmund Optics) and collected using a photomultiplier tube (model H10721-210; Hamamatsu). An amplifier (model C7319; Hamamatsu) was used to convert the current output from the photomultiplier tube to a voltage signal that was passed through a low-pass filter. The analog voltage signals were then digitized using an acquisition card (National Instruments). To minimize autofluorescence of the optical fiber, the recording fiber was photobleached using a high-power LED before recording. The excitation light power at the tip of the optical fiber was 20-30 μW and was delivered at 100 Hz with 5-ms pulse duration. Background autofluorescence was subtracted from the recorded signals in the subsequent analysis. A 635-nm laser (1-300 mW; LL-Laser, China) was used for optogenetic stimulation with the light power at the tip of the fiber set at 10 mW. Optical stimulation was delivered at 20 Hz (20-ms pulse duration) with a total of 1-20 pulses simultaneously with photometry recording. Where indicated, the mice received an intraperitoneal injection of SCH-23390 (2 mg/kg body weight).

When using the red-shifted GRAB_DA_ sensors, AAV-hsyn-rDA3m (300 nl) or AAV-hsyn-rDA3mut (300 nl) was injected into the CeA using the following coordinates: AP: −1 mm relative to Bregma; ML: ±2.5 mm relative to Bregma; and DV: −4.3 mm from the dura. In addition, AAV-hsyn-rDA2m (300 nl) or AAV-hsyn-rDA2mut (300 nl) was injected in the mPFC and NAc as described above, and AAV-hsyn-ChR2-YFP (300 nl) was injected into the VTA as described above. Two weeks after injection, photometry recording (FPS-410/470/561; Inper) was performed was described above. A 488-nm laser (1-160 mW, LL-Laser, China) was used for optogenetic stimulation, with the light power at the tip of the fiber set at 10 mW. Optical stimulation was delivered at 20 Hz (1, 5, or 10 s duration) simultaneously with photometry recording. Where indicated, the mice received an intraperitoneal injection of SCH-23390 (6 mg/kg body weight).

### Fiber photometry recording of DA signals in mice receiving water rewards

AAVs carrying GRABDA dopamine sensors were injected into the NAc of DAT-Cre mice (300 nL, unilateral injection; gGRAB_DA2m_: AAV9-hsyn-gDA2m, titer = 2.3×10^13^ vg ml^−1^; gGRAB_DA3m_: AAV9-hsyn-gDA3m, titer = 1.3×10^13^ vg ml^−1^). For red GRAB_DA_ sensors, we injected a mixture of AAVs carrying rGRAB_DA_ and GFP (3:1 mixture, 300nL total volume; GFP: AAV8-CAG-GFP, titer = 6.7 x 10^12^ vg ml^−1^; rGRAB_DA1m_: AAV9-hsyn-rDA1m, titer = 2.5×10^13^ vg ml^−1^; rGRAB_DA3m_: AAV9-hsyn-rDA3m, titer = 6×10^12^ vg ml^−1^). NAc coordinates were, from bregma: AP 1.5 mm, ML 1.7 mm, DV 4.0 mm, angled 4 degrees forward. Mice 1,2,5-10 also received injections of AAV8-hSyn-FLEX-ChrimsonR-tdTomato (titer = 3.9×10^12^ vg ml^−1^) in VTA and SNc (300 nL each, unilateral; coordinates, from bregma: VTA = AP −3.0 mm, ML 0.6 mm, DV 4.3 mm; SNc = AP −3.0 mm, ML 1.6 mm, DV 4.2 mm). Mice 3,4 received injections of AAV5-CAG-flex-tdTomato (titer = 4.8×10^12^) in VTA and SNc (300 nL each, unilateral). An optic fiber was implanted targeting NAc (400 um diameter, NA = 0.48, Doric lenses).

At least 2 weeks after surgery, mice were water restricted (to 85% of initial body weight) and trained to receive water rewards while head-fixed. After mice consistently licked to water delivery (typically requiring 3 days of training), we recorded dopamine sensor responses to unpredicted delivery of water droplets of various sizes (1, 2, 4, or 8 μL; inter-reward interval = 8-20 seconds, uniformly distributed; 60 trials per session, 15 trials per reward size, randomly interleaved; in some sessions, only 2 or 8 μL of water was given). Photometry signals were collected with a bundle-imaging fiber photometry system (Doric lenses). For green dopamine sensors, we used a blue LED (460-490 nm, 75 μW measured at tip of patch cord) to excite the sensor and the isosbestic wavelength as a control signal (410-420 nm LED, 60 μW). For red dopamine sensors, we used a yellow LED (555-570 nm, 110 μW) to excite the sensor and GFP signals as a control (460-490 nm LED, 75 μW). Imaging was performed at 20 Hz. Photometry data was processed offline as follows. ΔF/F_0_ = (F-F_0_)/F_0_ was computed by defining F_0_ as the 10^th^ percentile of each signal within a sliding 30 second window (excluding reward responses, defined as the 5 seconds following reward). Then, linear regression was performed between the sensor signal and the control signal (either isosbestic wavelength or GFP) in ITI periods (5 seconds following reward delivery excluded). The resulting predicted signal was subtracted from the sensor signal to produce the final de-noised signal.

### Two-photon *in vivo* imaging in mice

Adult (7-8 weeks of age) female C57BL/6N mice were used for these experiments. The mice were anesthetized with isoflurane (3% induction, followed by 1-1.5% maintenance), the skin and skull above the motor cortex were removed, and a metal recording chamber was affixed to the head. AAV expressing either gDA3m, gDA3h, dLight1.3b, or mEGFP (200 nl each, full titer) was then injected into the motor cortex using the following coordinates: AP: +1.0 mm relative to Bregma; ML: ±1.5 mm relative to Bregma; and DV: − 0.5 mm from the dura). A 4 mm x 4 mm square glass coverslip was then used to cover the opening in the skull. A stainless-steel head holder was attached to the skull to head-fix the animal’s head and reduce motion-induced artifacts during imaging. Two weeks after virus injection, the mice were habituated for approximately 10 min on the treadmill imaging apparatus to minimize stress. The motor cortex was imaged at a depth of 100-200 μm below the pial surface using Prairie View 5.5.64.100 software with an Ultima Investigator 2-photon microscope (Bruker) equipped with a 16×/0.80 NA water-immersion objective (Olympus) and an InSight X3 tunable laser (Spectra-Physics). A 920-nm laser was used for excitation, and a 525/70-nm emission filter was used to collect the fluorescence signal at a sampling rate of 1.5 Hz. For the forced running paradigm, running speed was set at 15 cm/s; for the tail shock paradigm, a 3-s electrical shock (0.7 mA) was delivered. For audio stimulation, a 1-s pulse of white noise (80 dB) was delivered. For image analysis, motion-related artifacts were corrected using the EZcalcium motion correction algorithm as described previously^63^. Fluorescence intensity measures at the ROIs was measured using ImageJ software. The fluorescent responses were calculated as [(F_raw_–F_baseline_)/F_baseline_], in which F_baseline_ was defined as the average fluorescence signal measured for 10 s prior to the behavior onset. The peak response during a behavior was calculated as the maximum *ΔF/F_0_*measured for 0-5 s after the behavior onset. The brain area was deemed responsive if the average response in a 5-s window surrounding the peak exceeded the sum of the baseline average and the baseline standard deviation. Hierarchical clustering was performed on the average of the fluorescence signals (forced running and shock) for each ROI. Euclidean distance and the Ward linkage metric were used after comparing multiple linkage metrics and clustering algorithms. Variations among individuals were minimized by normalizing the response to the maximum *ΔF/F_0_* across ROIs in a given mouse. The hierarchical method was used to reduce bias due to predetermining the cluster number.

### Quantification and statistical analysis

Except where indicated otherwise, all summary data are presented as the mean±SEM. Imaging data were processed using ImageJ (1.53c) or MATLAB software (matlab R2020a) and plotted using OriginPro 2020b (OriginLab), GraphPad Prism 8.0.2, or Adobe Illustrator CC. The change in fluorescence (*ΔF/F_0_*) was calculated using the formula [(F-F_0_)/F_0_], in which F_0_ is the baseline fluorescence signal. The SNR was calculated as the peak response divided by the standard deviation of the baseline fluorescence. Group differences were analyzed using a one-way analysis of variance (ANOVA) with Tukey’s multiple comparisons test, a one-way ANOVA with Dunnett’s multiple comparison test, a two-way ANOVA with Sidak’s multiple comparison test, or a two-tailed Student’s *t*-test (GraphPad Prism 8.0.2). Differences were considered significant at *p*<0.05; \**p*<0.05, \*\**p*<0.01, \*\*\**p*<0.001, \*\*\*\**p*<0.0001, and n.s., not significant (*p*>0.05). For all representative images and traces, similar results were obtained for >3 independent experiments.

## Data availability

The plasmids and sequences used to express the sensors in this study are available from Addgene. Source data will be provided upon reasonable request to the corresponding author.

## Code availability

The custom MATLAB codes, Arduino program, and ImageJ programs will be provided upon request to the corresponding author.

## Acknowledgments

The research was supported by the National Natural Science Foundation of China (grant nos. 31925017 and 31871087) and the NIH BRAIN Initiative (grant nos. 1U01NS113358 and 1U01NS120824); and by grants from the Feng Foundation of Biomedical Research, the Clement and Xinxin Foundation, the Peking-Tsinghua Center for Life Sciences, and the State Key Laboratory of Membrane Biology at Peking University School of Life Sciences to Y.L.; the NIH (grant no. R01DA004523) to J.T.W.; the NIH (grant no. R01MH125162) to M.W.-U.; the NIH (grant no. 5F32MH126505-02) to M.G.C. We thank I. Green and I. Tsutsui-Kimura for assistance for animal surgery. We thank X. Lei at PKU-CLS and the National Center for Protein Sciences at Peking University in Beijing, China, for support and assistance with the Opera Phenix high-content screening system and imaging platform.

## Author contributions

Y.L. supervised the study. Y. Zhuo. and Y.L. designed the study. Y. Zhuo., X.Y., Y.W., G.L., H.W. and Y. Zheng performed the experiments related to the development, optimization, and characterizing of the sensors in cultured HEK293T cells and in neurons. Y. Zhuo., R.C., and T.Q. performed the surgery and two-photon imaging experiments related to the validation of the sensors in acute brain slices. J.T.W. performed the characterization in acute brain slices containing the striatum or SNc. H.D., J.W., B. Li., and X.M. performed the *in vivo* fiber photometry recoding during optogenetic stimulation. M.G.C. performed the fiber photometry recording in the mouse NAc for the independent validation of *in vivo* sensor comparison under the supervision of M.W.-U. Y. Zhuo and B. Luo performed the *in vivo* fiber photometry recording in the NAc during mating behavior. B. Luo performed the *in vivo* fiber photometry recording during foot shock and the Pavlovian conditioning task with help from H.D. Y. Zhuo performed the *in vivo* two-photon imaging of the motor cortex. All authors contributed to the interpretation and analysis of the data. Y. Zhuo and Y.L. wrote the manuscript with contributions from all authors.

---

**Extended Data Fig. 1.**
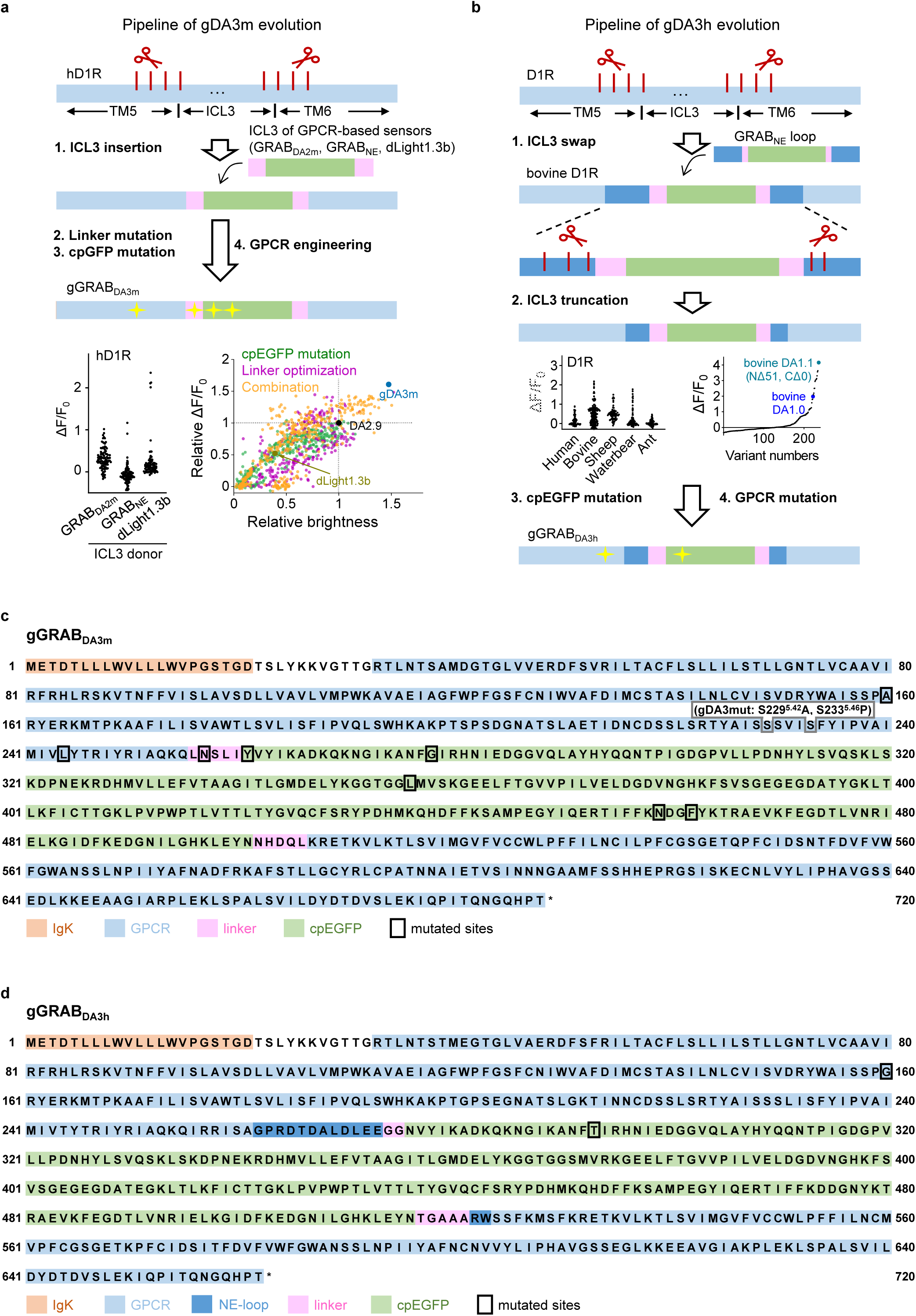
Strategy for optimizing and screening the green GRAB_DA_ sensors. **a**, A flowchart showing the development process (top) and screening (bottom) of the gDA3m sensor. ΔF/F_0_ represents the fluorescence change of sensor variants in response to 100 μM DA. The ICL3 domain of human D_1_R was replaced by the entire ICL3 (including linker and cpGFP) derived from several existing GPCR-based sensors (GRAB_DA2m_, GRAB_NE_ and dLight1.3b). Newly generated candidate with highest ΔF/F_0_ after ICL3 replacement (ICL3 from dLight1.3b) was then selected for further cpEGFP optimization, linker optimization and GPCR engineering. **b**, A flowchart showing the development process and screening of the gDA3h sensor. The ICL3 domains of dopamine D1 receptors from diverse species were replaced by the entire ICL3 from GRAB_NE_ sensor. Further optimization on the best chimera candidate (with bovine D_1_R backbone) includes ICL3 truncation for an optimal length, cpGFP mutation for improved brightness and response, as well as GPCR engineering for affinity tunning. **c**, Amino acids sequence of the gDA3m sensor. The mutations adopted in the gDA3m sensor are indicated by the black box. The serine residue at position 229^5.42^ in the human D_1_R was mutated to an alanine to generate the gDA3-mut sensor (indicated by the gray box). **d**, Amino acids sequence of the gDA3h sensor. The mutations adopted in the gDA3h sensor are indicated by the black box.

**Extended Data Fig. 2.**
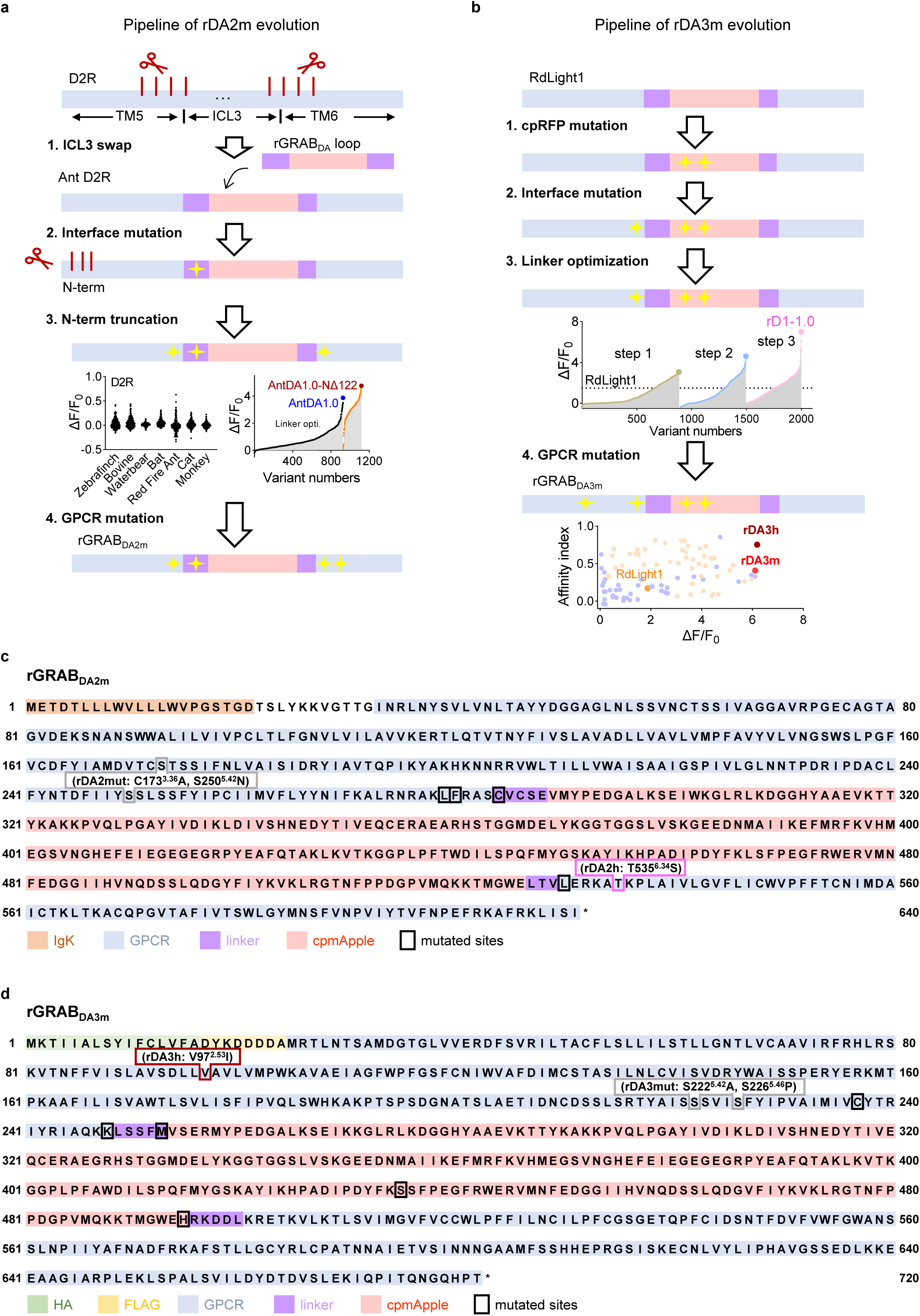
Strategy for optimizing and screening the green GRAB_DA_ sensors. **a**, A flowchart showing the development process and screening of the rDA2m sensor. The ICL3 domains of dopamine D2 receptors from diverse species were replaced by the entire ICL3 from rGRAB_DA_ sensor. Further optimization on the best chimera candidate (with ant D_2_R backbone) includes interface mutation (loop adjacent sites on the TM5 and TM6 of GPCR), receptor N-terminus truncation and GPCR mutation. **b**, A flowchart showing the development process and screening of the rDA3m and rDA3h sensor. The cpRFP module, RFP-GPCR interface, linker region and GPCR backbone of previously reported red dopamine sensor RdLight1 were systematically optimized. **c**, Amino acids sequence of the rDA2m sensor. The mutations adopted in the rDA2m sensor are indicated by the black box. The tyrosine residue at position 535^6.34^ in the ant D_2_R was mutated to serine to generate the high affinity rDA2h sensor (indicated by the magenta box). The cysteine to alanine mutation at position 173^3.36^ and serine to asparagine at position 250^5.42^ were adopted to generate the rDA2-mut sensor (indicated by the gray box). **d**, Amino acids sequence of the rDA3m sensor. The mutations adopted in the rDA3m are indicated by the black box. The valine residue at position 97^2.53^ in the human D_1_R was mutated to isoleucine to generate the high affinity rDA3h sensor (indicated by the dark red box). The serine to alanine mutation at position 222^5.42^ and serine to proline at position 226^5.46^ were adopted to generate the rDA3-mut sensor (indicated by the gray box).

**Extended Data Fig. 3.**
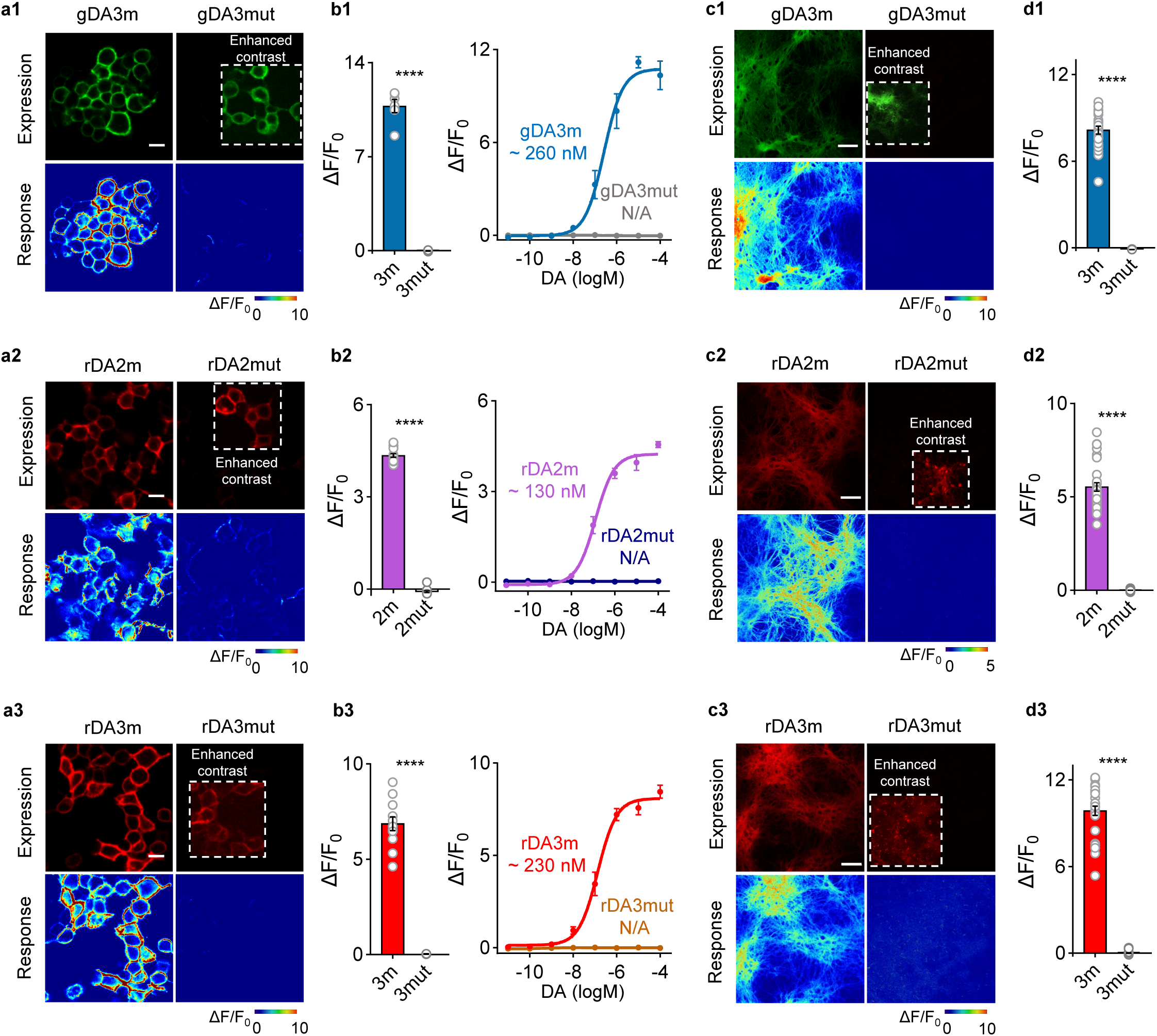
Performance of DA-insensitive mutant sensors. **a**, Representative images showing sensor expression (top) in HEK293T cells and fluorescence response to 100 μM DA (bottom) of indicated sensor variants. Scale bar, 20 μm. **b**, Group summary of maximal *ΔF/F_0_* in response to 100μM DA (left) and titration DA curves (right) of indicated sensors in HEK293T cells. Left, *n*=6, 6, 15, 15, 12, 3 wells for gDA3m, gDA3mut, rDA2m, rDA2mut, rDA3m and rDA3mut. Each well contains 400-500 cells. Two-tailed Student’s t-test was performed. Right, *n*=3 wells (with 400-500 cells per well) for each group. **c**, Representative images showing sensor expression (top) in cultured neurons and fluorescence response to 100 μM DA (bottom) of indicated sensor variants. Scale bar, 50 μm. **d**, Group summary of maximal *ΔF/F_0_* of indicated sensors in response to 100μM DA in cultured neurons. *n*=60 neurons from 4 cultures for rDA2mut, n=30/2 for others. Two-tailed Student’s t-test was performed.

**Extended Data Fig. 4.**
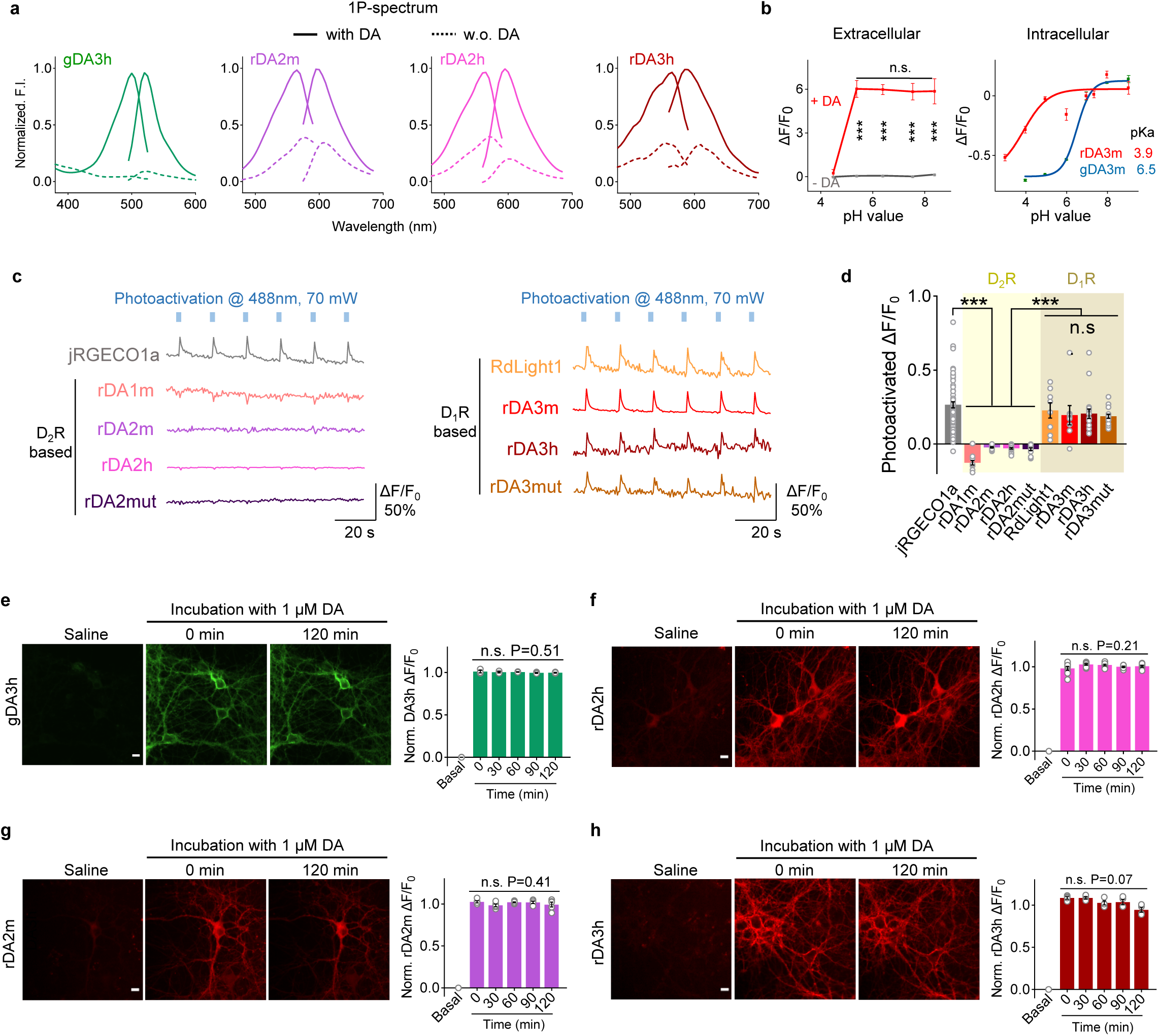
Characterization of GRAB_DA_ sensors in HEK393T cells and cultured neurons. a, Excitation and emission spectra for the indicated sensors in the absence and presence of DA. **b**, The effect of pH on GRAB_DA_ signals. Left, quantification for DA- or buffer-induced fluorescence responses of indicated sensors under different extracellular pH conditions. n=3 wells with 400-500 cells per well. Right, quantification for the relative buffer-induced fluorescence response of indicated sensors. The sensor-expressing HEK293T cells were gently permeabilized by detergent Triton-X100 (0.3% for ∼ 5 minutes). The fluorescence intensity of pH 6.95 was set as F_0_ and the relative fluorescence changes in each pH value were plotted. n=3 wells (with ∼2 − 10^5^ cells per well). **c-d**, Representative traces (**c**) and group summary of *ΔF/F_0_* (**d**) of indicated sensors upon blue-light illumination. *n*= 18, 14, 18, 16 cells for rDA2h, rDA2mut, rDA3h and rDA3mut; other data replotted from Fig. 2g. **e-h**, Representative images (left) and quantification (right) of the change in sensor fluorescence in response to 2-h application of 100μM DA. Scale bar, 20 μm. *n*=3, 8, 5, 4 cultures for gDA3h, rDA2h, rDA2m and rDA3h. One-way ANOVA test was performed for DA-application groups. n.s. p=0.5104, 0.2183, 0.4101, 0.0652 for gDA3h, rDA2h, rDA2m and rDA3h.

**Extended Data Fig. 5.**
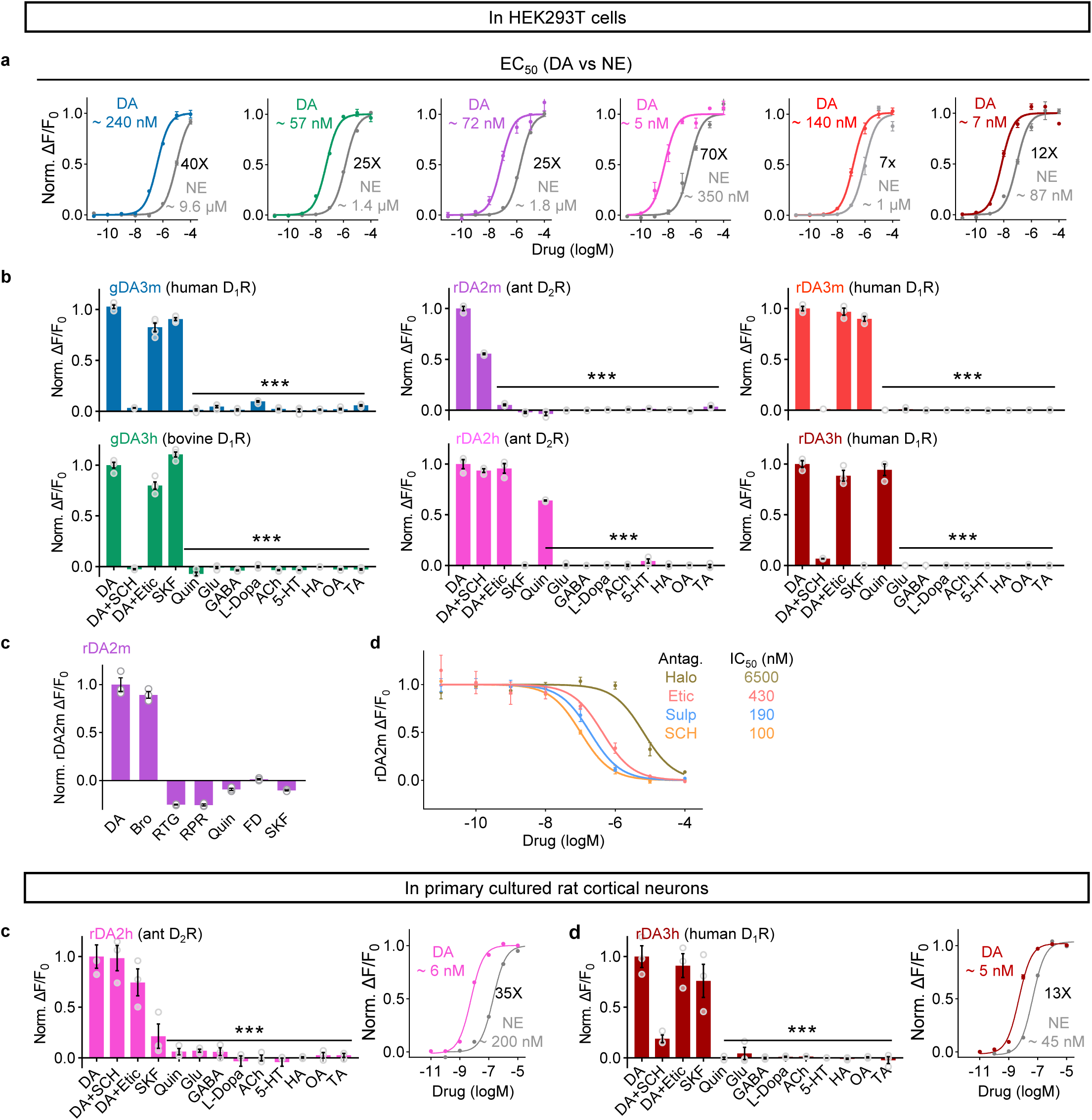
Pharmacological profiles of new GRAB_DA_ sensors measured in cultured cells. **a**, Titration curves of indicated sensors for the response to DA or NE in HEK293T cells. *n*=3 cells with 400-500 cells per well. **b**, The normalized *ΔF/F_0_* in sensor-expressing HEK293T cells in response to the indicated compounds. Antagonists were applied at 10μM, others at 1μM. *n*=4 wells for gDA3m and gDA3h, *n*=3 wells for others. **c**, The normalized *ΔF/F_0_* in rDA2m-expressing HEK293T cells in response to indicated DA agonists. Bromocriptine (Bro), Rotigotine (RTG), D_2_R/D_1_R agonists; Ropinirole (RPR), Quin, D_2_R-specific agonists; Fenodopam (FD), SKF, D_1_R-specific agonist. All chemicals were bath-applied in 100 μM. One-way Anova, post hoc Dunnett’s test was performed. n.s. p=0.1074 between DA and Bro. **d**, Titration curves of indicated dopamine receptor antagonists. The fluorescence intensity in the presence with 10 μM DA was set as F_0_ and the relative fluorescence changes under indicated compound concentration were plotted. **e-f**, Pharmacological specificity (left) and titration curves of indicated sensors for the response to DA or NE (right) in cultured neurons. Left, antagonists were applied at 10μM, others at 1μM. *n*=3 wells. One-way Anova, post hoc Dunnett’s test was performed. rDA2h, n.s. p=0.9998, 0.1458 between DA and DA+SCH, or DA+Etic; rDA3m, n.s. p=0.9591, 0.1309 between DA and DA+Etic, or SKF.

**Extended Data Fig. 6.**
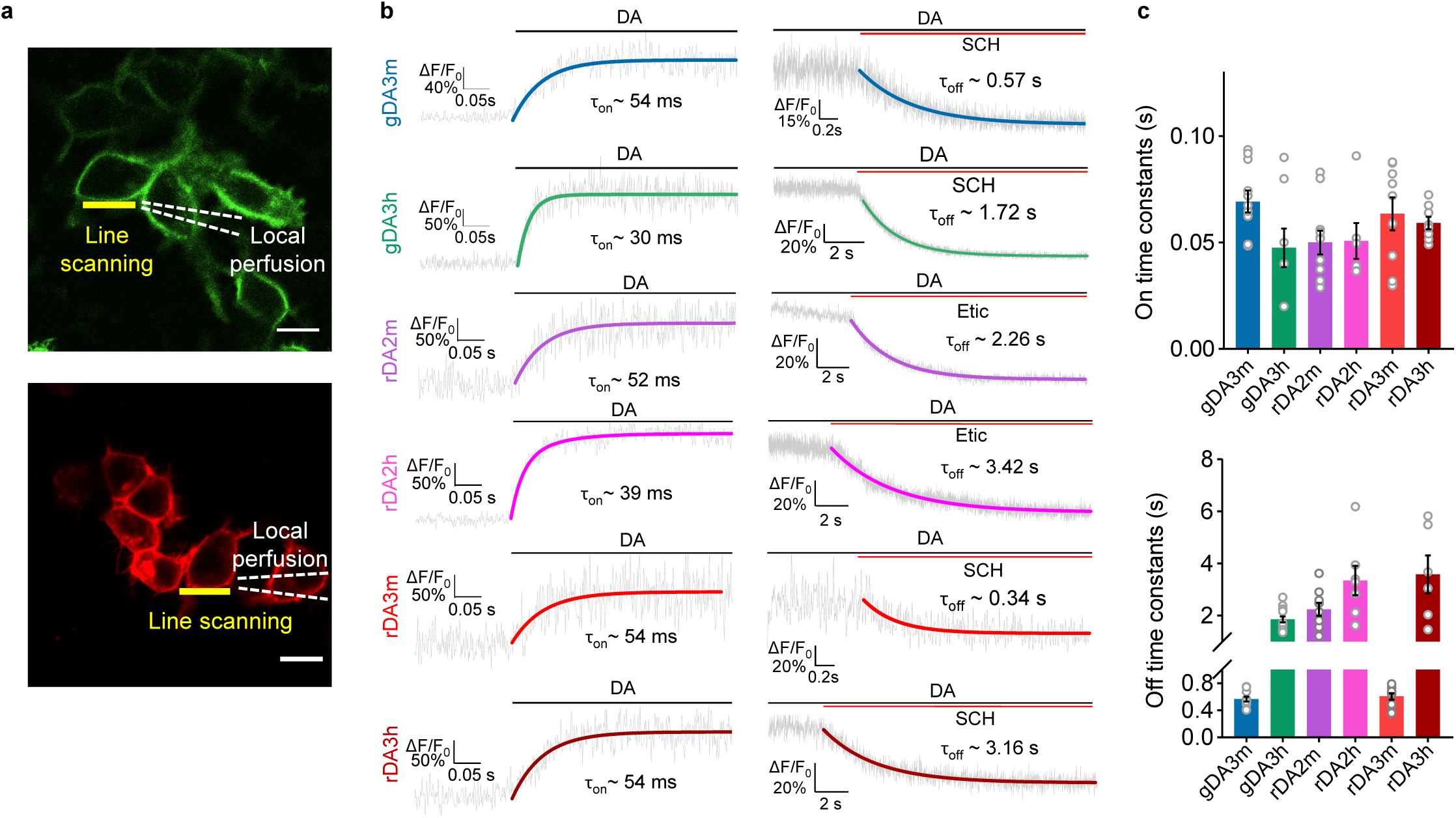
Kinetics measurement of new GRAB_DA_ sensors in HEK293T cells. **a**, Schematic illustration showing the local perfusion system using a glass pipette containing 100 μM DA and/or receptor-specific antagonist positioned above the sensor-expressing cell. The yellow line indicates the area for line scanning. The dash lines indicate the pipette. Scale bar, 20 μm. **b**, Representative traces showing the response measured using line-scanning; when indicated, DA and receptor-specific antagonist were puffed onto the cell. The trace were the average of 3 different ROIs on the scanning line. Data are shown as mean±SD. Each trace was fitted with a single-exponential function to determine the τ_on_ (left) and τ_off_ (right). **c**, Group summary of τ_on_ and τ_off_. τ_on_, *n*= 11, 8, 11, 6, 9, 8 cells for gDA3m, gDA3h, rDA2m, rDA2h, rDA3m, rDA3h. τ_off_, *n*=10, 14, 9, 7, 10, 6 cells for gDA3m, gDA3h, rDA2m, rDA2h, rDA3m, rDA3h.

**Supplementary Table 1:** Properties of genetically encoded GPCR-based dopamine sensors.

| Sensor | Color | GPCR backbone | Maximal brightness | Response (maximum $\Delta F/F_0$) | Apparent affinity ($EC_{50}$, nM) | On kinetics ($\tau$, ms) | Off kinetics ($\tau$, s) | 1-photon Ex./Em. (nm) | Blue-light photoactivated $\Delta F/F_0$ | Source |
| --- | --- | --- | --- | --- | --- | --- | --- | --- | --- | --- |
| gDA2m | G | D<sub>2</sub>R (human) | 2.4<sup>a,c</sup> | 2.4<sup>a</sup><br>3.3<sup>b</sup> | 60<sup>a</sup><br>45<sup>b</sup> | 60* | 0.71* | ND | - | ref. <sup>16</sup> |
| dLight1.3b | G | D<sub>1</sub>R (human) | 1.4<sup>a,c</sup> | 3.7<sup>a</sup><br>5.2<sup>b</sup> | 870<sup>a</sup><br>690<sup>b</sup> | ND | ND | ND | - | ref. <sup>13</sup> |
| gDA3m | G | D<sub>1</sub>R (human) | 3.0<sup>a,c</sup> | 10.0<sup>a</sup><br>24.0<sup>b</sup> | 89<sup>a</sup><br>120<sup>b</sup> | 69 | 0.56 | 495/520 | - | this paper |
| gDA3h | G | D<sub>1</sub>R (bovine) | 3.5<sup>a,c</sup> | 12.4<sup>a</sup><br>13.4<sup>b</sup> | 22<sup>a</sup><br>12<sup>b</sup> | 48 | 1.85 | 500/520 | - | this paper |
| rDA1m | R | D<sub>2</sub>R (human) | 2.0<sup>a,d</sup> | 1.2<sup>a</sup><br>1.5<sup>b</sup> | 370<sup>a</sup><br>100<sup>b</sup> | 80* | 0.77* | 565/595* | -0.12 | ref. <sup>16</sup> |
| rDA2m | R | D<sub>2</sub>R (red fire ant) | 8.2<sup>a,d</sup> | 5.3<sup>a</sup><br>6.6<sup>b</sup> | 210<sup>a</sup><br>140<sup>b</sup> | 50 | 2.24 | 565/595 | -0.02 | this paper |
| rDA2h | R | D<sub>2</sub>R (red fire ant) | 8.7<sup>a,d</sup> | 2.4<sup>a</sup><br>3.3<sup>b</sup> | 9.8<sup>a</sup><br>6.0<sup>b</sup> | 50 | 3.35 | 565/595 | -0.03 | this paper |
| RdLight1 | R | D<sub>1</sub>R (human) | 1.0<sup>a,d</sup> | 4.4<sup>a</sup><br>2.6<sup>b</sup> | 2700<sup>a</sup><br>310<sup>b</sup> | ND | ND | 560/588* | 0.23 | ref. <sup>15</sup> |
| rDA3m | R | D<sub>1</sub>R (human) | 4.3<sup>a,d</sup> | 14.6<sup>a</sup><br>12.6<sup>b</sup> | 140<sup>a</sup><br>40<sup>b</sup> | 64 | 0.61 | 560/595 | 0.19 | this paper |
| rDA3h | R | D<sub>1</sub>R (human) | 3.8<sup>a,d</sup> | 14.2<sup>a</sup><br>10.1<sup>b</sup> | 22<sup>a</sup><br>5.5<sup>b</sup> | 60 | 3.60 | 565/585 | 0.20 | this paper |

G: green; R: red; Ex: excitation wavelength; Em: emission wavelength; ND: not determined; <sup>a</sup> determined in HEK293T cells; <sup>b</sup> determined in cultured neurons; <sup>c</sup> relative to basal brightness of gDA2m; <sup>d</sup> relative to basal brightness of rDA1m; \* previously reported results; kinetics was estimated in HEK293T cells to puff ligand application; photoactivation was estimated in HEK293T cells with 488-nm illumination in ligand-free condition.
**Note:** All data were collected in this paper unless indicated.

**Extended Data Fig. 7.**
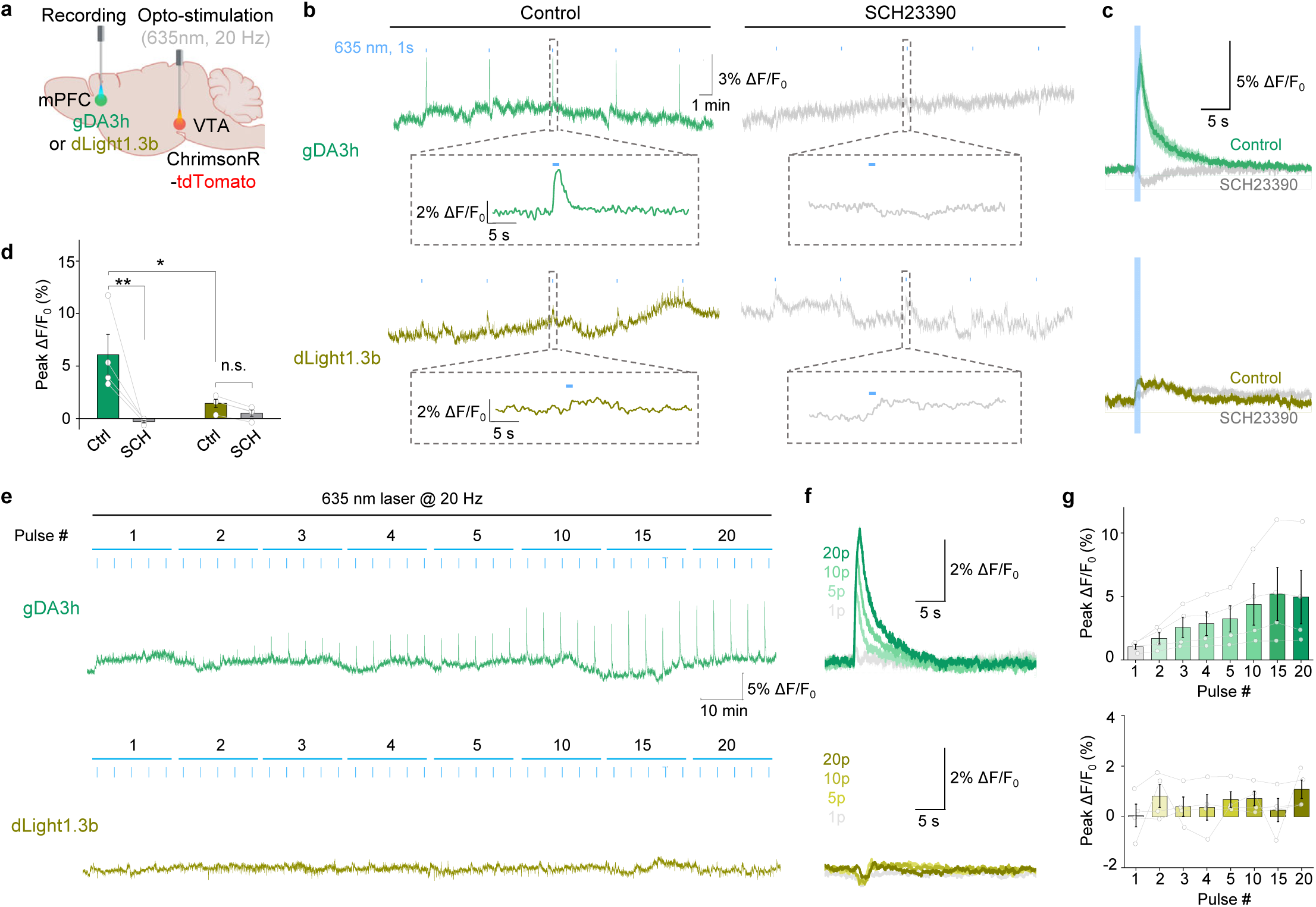
gGRAB_DA3h_ sensors report optogenetically-elicited DA release in the mouse mPFC a, Schematic illustration depicting the experimental design for panel **b-g**. **b**, Representative fluorescence changes and zoom-in view (indicated by dashed box) of indicated sensors during optogenetic stimulations under control condition or in the presence of SCH-23390 (SCH). **c**, Average traces of the change in gDA3h (top) or dLight1.3b (bottom) fluorescence from a mouse. Data are shown as mean±SD. **d**, Group summary of *ΔF/F_0_* for the indicated sensors. *n*=4 mice for gDA3h and dLight1.3b, respectively. One-way ANOVA, post hoc Tukey’s test was performed. **p=0.0035 for gDA3h; n.s. p=0.9122 for dLight1.3b; *p=0.0295 between gDA3h and dLight1.3b. **e-f**. Example fluorescence response (**e**) and corresponding average traces (**f**) of gDA3h (top) or dLight1.3b (bottom) to indicated optogenetic stimulation. The average traces are shown as mean±SD. **g**, Group summary of peak *ΔF/F_0_* of gDA3h or dLight1.3b in response to indicated optogenetic stimulation. *n*=4 mice for gDA3h and dLight1.3b.

**Extended Data Fig. 8.**
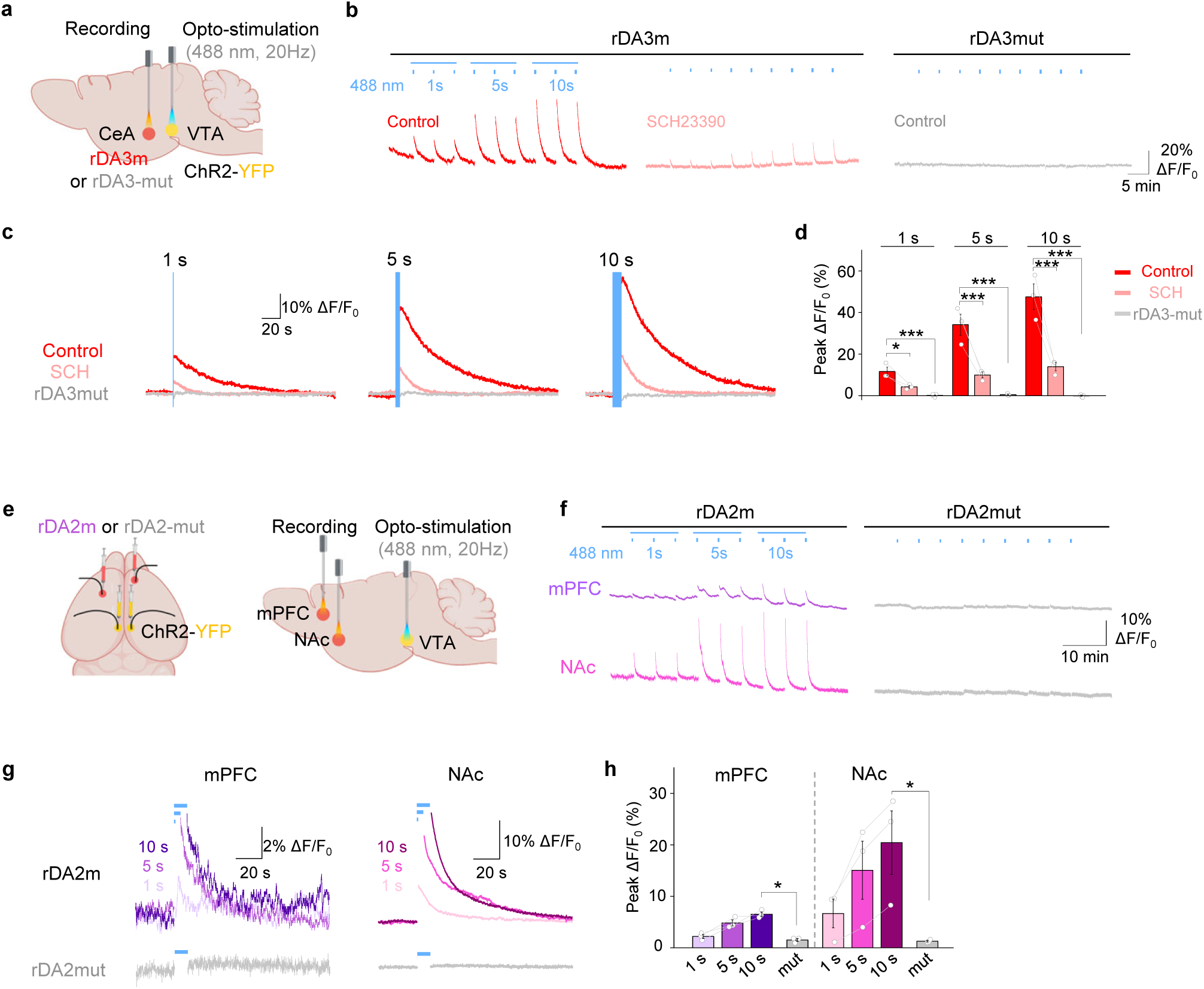
rGRAB sensors report optogenetically-elicited DA release in multiple brain regions *in vivo*. **a**, Schematic illustration depicting the experimental design for panel **b-d**. **b**, Representative traces of rDA3m or rDA3mut signals during optogenetic stimulations. rDA3m signals were measured before and after SCH-23390 (SCH) administration. **c**, Average traces of the change in sensor fluorescence to 1-, 5- or 10-s opto-stimulation from a mouse. Data are shown as mean±SD. The blue shaded area indicates the application of opto-stimulation. **d**, Group summary of peak response of rDA3m or rDA3mut to indicated optogenetic stimulation. *n*=3 mice for rDA3m and *n*=5 for rDA3mut. Two-tailed Student’s t-test was performed. p=0.0278, 0.0101, 0.0068 between control and SCH to 1-, 5-, 10-s opto-stimulation. p=0.0003, 0.0001, <0.0001 between rDA3m and rDA3mut to 1-, 5-, 10-s opto-stimulation. **e**, Schematic illustration depicting the experimental design for panel **e-h**. **f**, Representative traces of sensor signals simultaneously recorded in the mPFC (top) and NAc (bottom) during optogenetic stimulations. **g**, Average traces of the change in sensor fluorescence in the mPFC (left) and NAc (right) to indicated optogenetic stimulation from a mouse. Data are shown as mean±SD. The length of blue lines indicates the duration of opto-stimulation. **h**, Group summary of peak response of rDA2m or rDA2mut to indicated optogenetic stimulation. *n*=3 mice for rDA2m and rDA2mut. Two-tailed Student’s t-test was performed between rDA3m and rDA3mut response upon 10-s opto-stimulation. p=0.0007 for mPFC, p=0.0364 for NAc.

**Extended data Fig. 9.**
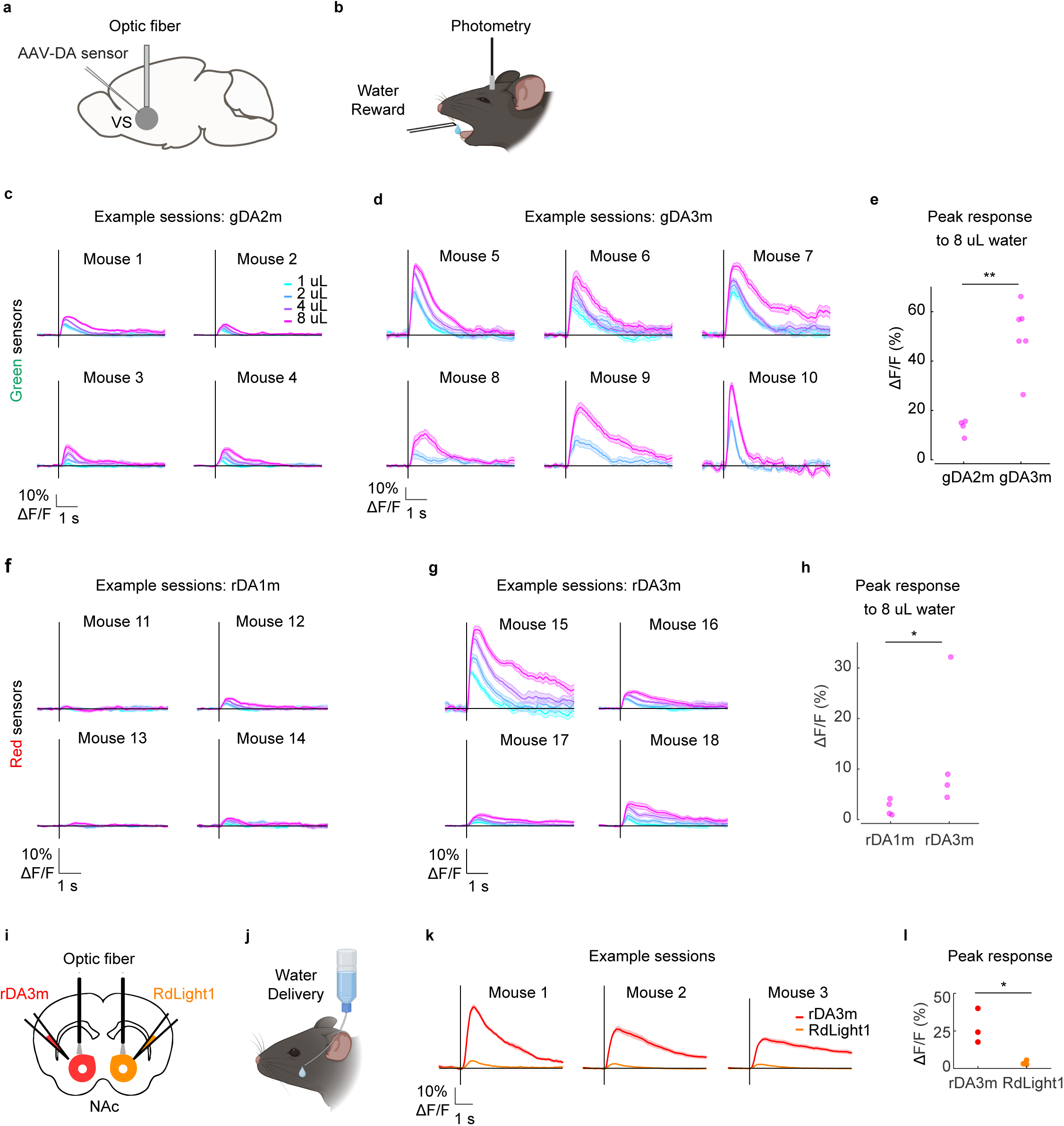
*In vivo* comparison of the third-generation DA sensors versus previous variants in water-restricted mice receiving water rewards. **a**, Diagram of mouse surgical procedure. AAVs carrying gGRAB_DA2m_, gGRAB_DA3m_, rGRAB_DA1m_, or rGRAB_DA3m_ were injected unilaterally into NAc. An optic fiber was implanted above the injection site. **b**, Illustration of behavioral experiment. **c**, Recording sessions from gDA2m mice. Vertical black bars indicate water delivery. Colors indicate water volume. **d**, Recording sessions from gDA3m mice. **e**, Peak response to 8 μL water for the sessions shown in **c** and **d**. ** p = 0.0095, Mann-Whitney U test. **f**, Recording sessions from rDA1m mice. **g**, Recording sessions from rDA3m mice. **h**, Peak response to 8 μL water for the sessions shown in **f** and **g**. ** p = 0.0286, Mann-Whitney U test. **i-j**, Schematic illustration depicting the mouse surgical procedure and the experimental design for panel **k-l**. **k**, Recording sessions from 3 mice. Vertical black bars indicate water delivery. Colors indicate sensor version. **l**, Peak response of rDA3m and RdLight1 for the sessions shown in **k**. * p = 0.0249, Two-tailed Student’s t-test.

**Extended data Fig. 10.**
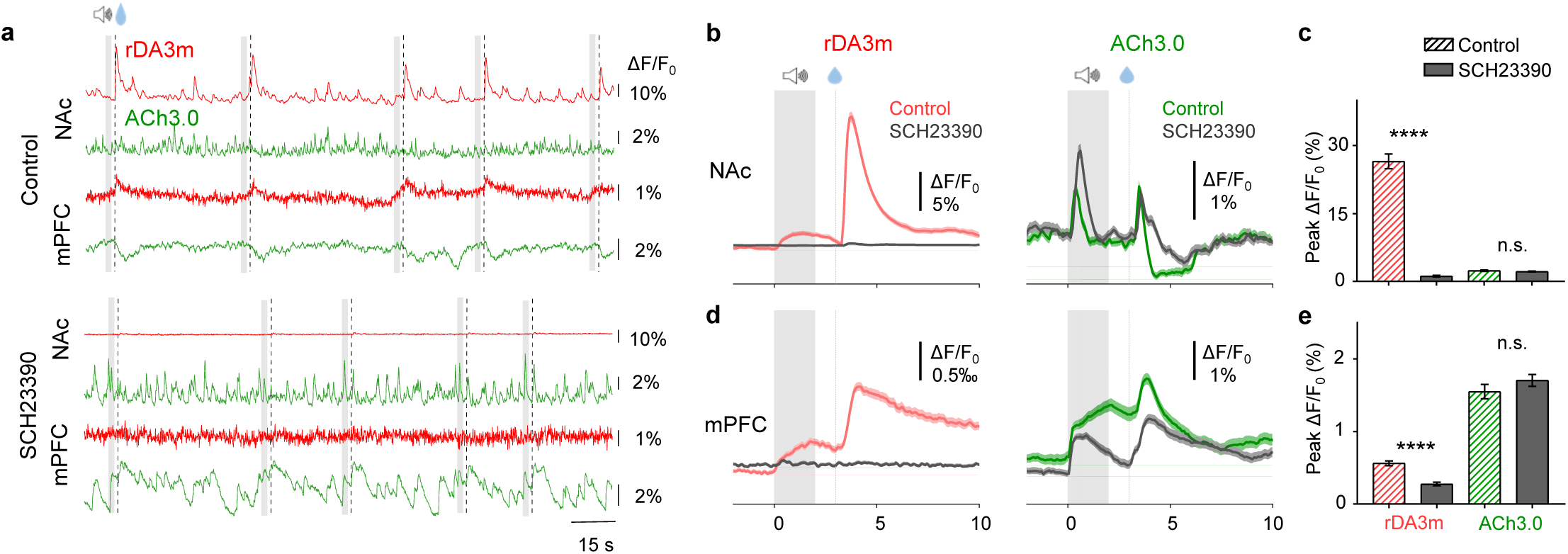
The signals in the mouse NAc and mPFC during Pavlovian conditioning. **a**, Representative fluorescence signals recorded during consecutive water trials pre (top, control) and post SCH-23390 (bottom, SCH-23390) treatment. The audio and water delivery are indicated above. **b**, Averaged traces of rDA3m (left) and ACh3.0 (right) fluorescence measured in the NAc from a mouse under control condition or in the presence of SCH-23390. Shown are more than 50 consecutive trials (mean±SD) in one mouse. The grey shaded area indicates the application of audio. The dashed line indicates the delivery of water. **c**, Group summary of the peak fluorescence change of rDA3m and ACh3.0 signals in the NAc under the indicated condition. *n*= 155 trials from 3 mice for each group. Two-tailed Student’s t-test was performed between control and SCH-23390 group. p=0.2624 for ACh3.0. **d-e**. same as (**b-c**) with simultaneously recorded rDA3m and ACh3.0 signals in the mPFC. Two-tailed Student’s t-test was performed between control and SCH-23390 group. p=0.2274 for ACh3.0.

**Extended data Fig. 11.**
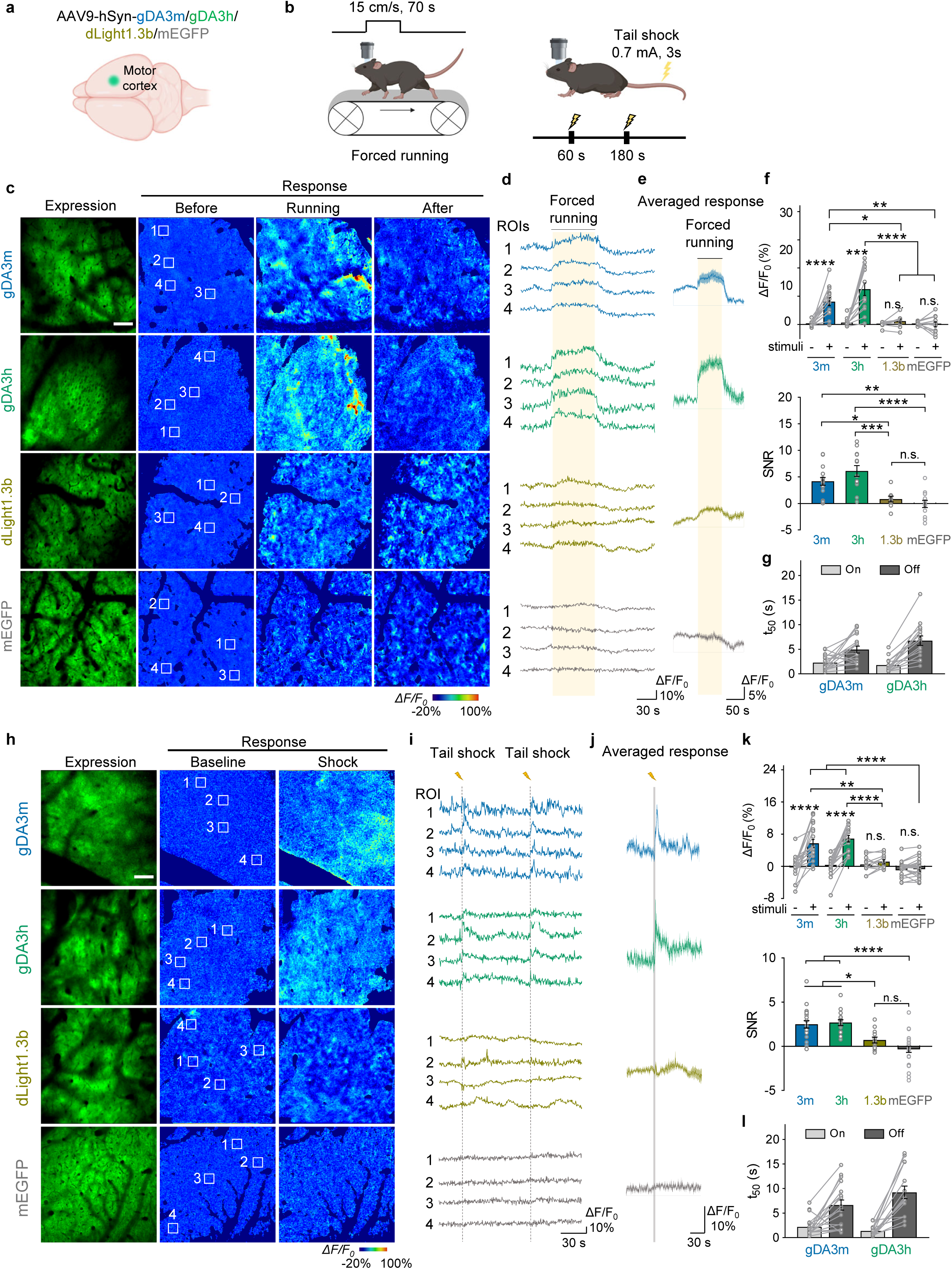
*In vivo* two-photon imaging of cortical DA dynamics in mice. **a-b**, Schematic illustration depicting the experimental design for panel **c-j**. **c-e**, Representative expression and pseudocolored response images (**c**), representative traces measured at the indicated ROIs (**d**), and average traces per forced running (**e**) measured in the head-fixed mice expressing gDA3m, gDA3h, dLight1.3b or mEGFP. Scale bar, 100 μm. **f**, Group summary of the peak fluorescence response (top) and SNR (bottom) measured during forced running in the motor cortex of mice expressing gDA3m, gDA3h, dLight1.3b and mEGFP. n=14 trials from 4 mice (15/4) for gDA3m, 13/4 for gDA3h, 9/3 for dLight1.3b, 12/4 for mEGFP. Paired two-tailed Student’s t-test was performed within group. One-way ANOVA, post hoc Tukey’s test was performed across sensor groups. Response, p<0.0001 for gDA3m; p=0.0002, 0.0683, 0.6275 for gDA3h, dLight1.3b, mEGFP; p<0.0001 between gDA3h and dLight1.3b, or EGFP; p=0.0214 between gDA3m and dLight1.3b; p=0.0022 between gDA3m and mEGFP; p=0.1611 between gDA3m and gDA3h; p=0.9577 between dLight1.3b and mEGFP. SNR, p<0.0001 between gDA3h and mEGFP; p=0.0004 between gDA3h and dLight1.3b; p=0.0016 between gDA3m and mEGFP; p=0.0337 between gDA3m and dLight1.3b; p=0.8812 between dLight1.3b and mEGFP. **g**, Summary of the rise and decay t_50_ values (where applicable) of the gDA3m and gDA3h signals in response to forced running. **h-j**, Same as (**c-e**) except mice were subjected to tail shock. **k**, Group summary of the peak fluorescence response (top) and SNR (bottom) measured upon tail shock in the motor cortex of mice expressing gDA3m, gDA3h, dLight1.3b and mEGFP. n=19/4 for gDA3m, 16/4 for gDA3h, 12/3 for dLight1.3b, 26/4 for mEGFP. Paired two-tailed Student’s t-test was performed within group. One-way ANOVA, post hoc Tukey’s test was performed across sensor groups. Response, p<0.0001 for gDA3m and gDA3h; p=0.1774, 0.2524 for dLight1.3b, mEGFP; p<0.0001 between mEGFP and gDA3m, or gDA3h; p<0.0001 between gDA3h and dLight1.3b; p=0.0.0013 between gDA3m and dLight1.3b; p=0.7169 between gDA3m and gDA3h; p=0.3714 between dLight1.3b and mEGFP. SNR, p<0.0001 between mEGFP and gDA3m, or gDA3h; p=0.0186, 0.0104 between dLight1.3b and gDA3m, or gDA3h; p=0.2607 between dLight1.3b and mEGFP. **l**, Summary of the rise and decay t_50_ values of the gDA3m and gDA3h signals in response to tail shock. mEGFP data replotted from Fig. 6f.

